# Fold-and-fuse neurulation in zebrafish requires Vangl2

**DOI:** 10.1101/2023.11.09.566412

**Authors:** Jacalyn MacGowan, Mara Cardenas, Margot Kossmann Williams

## Abstract

Shaping of the future brain and spinal cord during neurulation is an essential component of early vertebrate development. In amniote embryos, primary neurulation occurs through a “fold-and-fuse” mechanism by which the edges of the neural plate fuse into the hollow neural tube. Failure of neural fold fusion results in neural tube defects (NTDs), which are among the most devastating and common congenital anomalies worldwide. Unlike amniotes, the zebrafish neural tube develops largely via formation of a solid neural keel that later cavitates to form a midline lumen. Although many aspects of primary neurulation are conserved in zebrafish, including neural fold zippering, it was not clear how well these events resemble analogous processes in amniote embryos. Here, we demonstrate that despite outward differences, zebrafish anterior neurulation closely resembles that of mammals. For the first time in zebrafish embryos, we directly observe enclosure of a lumen by the bilateral neural folds, which fuse by zippering between at least two distinct closure sites. Both the apical constriction that elevates the neural folds and the zippering that fuses them coincide with apical Myosin enrichment. We further show that embryos lacking *vangl2*, a core planar cell polarity and NTD risk gene, exhibit delayed and abnormal neural fold fusion that fails to enclose a lumen. These defects can also be observed in fixed embryos, enabling their detection without live imaging. Together, our data provide direct evidence for fold-and-fuse neurulation in zebrafish and its disruption upon loss of an NTD risk gene, highlighting the deep conservation of primary neurulation across vertebrates.

**Highlights:**

- The anterior neural tube of zebrafish undergoes “fold-and-fuse” neurulation to enclose a lumen, highlighting conservation of primary neurulation mechanisms across vertebrates.
- Anterior neural tube closure is delayed and abnormal in zebrafish embryos lacking the planar cell polarity gene *vangl2*, occurring by excessive “buttoning” rather than smooth “zippering” and failing to enclose a lumen.
- Neural tube defects (NTDs) are visible in fixed *vangl2* deficient embryos, enabling simple assessment of neural tube phenotypes with potential utility in screening NTD risk genes.

## Introduction

Neural tube defects (NTDs) such as spina bifida and anencephaly are among the most common and devastating congenital anomalies, affecting approximately 1 in 1,000 births in the United States (*1*) and even more worldwide (*2*). These conditions result from incomplete closure of the neural tube during embryogenesis, often leaving the neural tube lumen open to the outside of the body. Primary neurulation, which forms the neural tube within the head and trunk, is well characterized in amniote embryos like mouse, chick, and to a lesser extent, human. In these species, neural tube formation is driven by convergent extension (CE) of the developing neural plate followed by formation of hinge points (*3–10*) that elevate the bilateral neural folds and bend them toward each other (*9, 11, 12*). The neural folds then meet at the dorsal midline and fuse by zippering between discrete closure points (*13, 14*), completing the “fold-and-fuse” process that encloses the neural tube lumen. By contrast, the zebrafish spinal cord develops from the neural keel, a solid structure that later undergoes cavitation to form a central lumen (*15, 16*). The site of this lumen is established through a series of midline-crossing mitoses termed “C-divisions” which distribute one daughter cell of each side of the neural keel midline (*15, 17–22*). For this reason, neurulation in zebrafish has been likened to secondary neurulation in amniote embryos (*16, 23, 24*), during which the post-anal neural tube forms by condensation of mesenchymal cells (*25, 26*). Primary neurulation in zebrafish was therefore often viewed as fundamentally different from other vertebrates, making it a poor model for NTDs.

However, many studies have revealed that despite these outward differences, several hallmarks of primary neurulation are conserved in zebrafish. For example, CE morphogenesis narrows the neural plate (*21, 27*), and apical constriction at the midline forms a medial hinge point-like structure (*28, 29*). The neural folds were also shown to zipper closed in the forebrain region of zebrafish (*29*) in a fashion strikingly similar to mice (*13, 14*). These conserved neurulation mechanisms open the possibility of modeling NTDs, or aspects thereof, in the experimentally tractable zebrafish model. Indeed, previous studies have proposed bifurcation of pineal gland precursors and/or the dorsal roof plate as proxies for NTDs in zebrafish (*30–33*). These phenotypes are suggestive of reduced neural fold convergence, but it is unclear whether they exhibit – or whether zebrafish are capable of exhibiting – the open neural tubes that define NTDs.

This is illustrated by neural phenotypes in zebrafish embryos lacking the core planar cell polarity (PCP) gene *vangl2*. PCP signaling is a highly conserved regulator of vertebrate gastrulation and neurulation, and loss of PCP components in mouse, chick, and *Xenopus* disrupts CE of the neural plate, hinge point formation, and ultimately neural tube closure (*3, 4, 34–47*). Mutations in PCP genes (including *VANGL2*) are also associated with NTDs in several patient cohorts (*48–54*). Loss of PCP signaling similarly disrupts CE and neural tube development in zebrafish (*21, 27, 55–64*), but these phenotypes differ from those of amniotes. Reduced convergence of the neural plate in maternal-zygotic (MZ)*vangl2*/*trilobite (tri)-/-* zebrafish causes C-divisions to occur at lateral rather than midline positions of the future spinal cord, giving rise to ectopic bilateral lumens (*21, 22, 55*). While striking, these phenotypes do not outwardly resemble NTDs in other vertebrates, presumably because of the distinct mechanisms of neurulation and/or lumen formation between species.

Here, we carefully reevaluate neural tube development in wild-type (WT) and *vangl2* deficient zebrafish embryos with a focus on the future forebrain at early stages of neurulation. Live time-lapse imaging of neural tube closure revealed that, in addition to the previously described zippering of the anterior neural folds, WT zebrafish exhibit a distinct posterior site of neural fold fusion. The anterior and posterior openings zipper in opposite directions from a central point of contact to close the anterior neural tube. Using optical transverse sectioning of live embryos, we further showed that the bilateral neural folds fuse to enclose a lumen in a process strikingly similar to amniote neurulation. Neural fold fusion was delayed and neural groove formation was abnormal in *vangl2* deficient embryos, resulting in impaired midline convergence of pineal precursors and pit-shaped openings in the forebrain region that were readily visible in live and fixed embryos. Together, these data provide direct evidence for fold-and-fuse neurulation in zebrafish and a requirement for PCP-dependent CE in this process, demonstrating deep conservation of neurulation mechanisms among vertebrates.

## Results

### The zebrafish forebrain neural tube forms distinct anterior and posterior openings

A recent study used live time-lapse imaging to directly observe closure of the forebrain neural tube in zebrafish, revealing zippering of an eye-shaped opening between the fusing neural folds beginning at approximately 6-7 somite stage (*29*). To characterize neural tube closure more fully in WT embryos, we performed confocal time-lapse imaging of the anterior neural plate beginning at the 3 somite stage. In each of the 30 control embryos imaged, we observed the presence and zippering of an eye-shaped opening in the forebrain region as previously reported (*29*). However, examining the neural plate at earlier stages revealed a sequence of preceding morphogenetic changes (**Fig. 1**). At around the 5 somite stage, we observed a continuous keyhole-shaped groove in the neural plate midline with the round portion positioned anteriorly (**Fig. 1A-B**, green shading). The bilateral neural folds elevated on either side of this groove (**Supp. video 1**) and came together near the center of the keyhole to “pinch off” anterior and posterior portions that each zippered closed away from the “pinch point” (**Fig. 1A-B** and **Supp. Fig. 3**, white arrows, **Supp. video 2**). Elevation of the neural folds was accompanied by apical constriction of midline neuroectoderm cells (**Fig. 1D ’**, **Supp. Video 1**), which was previously reported and extensively quantified in the future forebrain and hindbrain (*28, 29*).

We next used an Sf9-mNeon intrabody (*65, 66*) to observe non-muscle Myosin II localization during these cell shape changes in the neural plate. Shortly before formation of the keyhole-shaped groove, Myosin accumulated in patches at the medial apical cortex of midline neuroectoderm cells as they constricted (**Fig. 1C-D, Supp video 3**). This is similar to reports in other apically constricting tissues like the invaginating mesoderm of *Drosophila* embryos (*67*) and hindbrain neural plate in zebrafish (*28*), and consistent with apical localization of phospho-Myosin light chain in medial hinge point of the zebrafish forebrain (*29*). Apical Myosin was first apparent in the anterior portion of the neural groove as it invaginated to form a pit (**Fig. 1C**, yellow arrows), which continued to deepen into the round end of the keyhole-shaped groove. Shortly after apical Myosin became visible anteriorly, we observed medial-apical patches within midline cells of the posterior neural groove (**Fig.1D**, white arrows). The apices of these cells constricted primarily in the ML dimension to create AP-elongated apical surfaces (**Fig.1D’**, white arrows), which was also observed during simultaneous invagination and convergence in *Drosophila* mesoderm, chicken neural tube, and (to a lesser degree) zebrafish hindbrain (*28, 45, 68, 69*). Previous studies showed that inhibiting Myosin contractility with Blebbistatin prevented medial hinge point formation and neural fold fusion in zebrafish (*28, 29*). This suggests that Myosin-driven constriction and invagination at the midline bring the two sides of the neural plate together to create the pinch point from which bidirectional neural fold fusion proceeds.

The posterior opening zippered closed first, beginning at the pinch point and continuing posteriorly (**Fig. 1A-B**, blue shading, **Supp. videos 1-4**), leaving the anterior portion to form the eye-shaped opening later. In some control embryos, a small opening at the posterior end of this zipper could later be seen completing closure. Once the posterior opening had zippered (mostly) closed, the neural folds of the anterior portion formed the posterior closure point of the eye-shaped opening (**Fig. 1A-B** and **Supp. Fig. 3**, yellow shading and arrows), as previously described (*29*). The anterior closure point of the eye-shaped opening arose from the anterior-most edge of the initial keyhole-shaped groove (**Fig. 1** and **Supp. Fig. 3**, yellow arrows), and the opening zippered closed predominantly from anterior to posterior (**Supp. video 5**). The posterior-most end of the posterior opening also completed its zippering at this stage. Notably, our live imaging also enabled examination of the relationship between midline-crossing C-divisions and neural tube closure. In embryos in which the left and right sides of the developing neural keel exhibited distinct levels of fluorescent protein expression (**Supp. video 2**), cells were only seen crossing the midline after the anterior eye-shaped opening had closed. This provides evidence that neural fold fusion precedes C-divisions.

Together, these observations delineate a complex series of morphogenetic events that close the anterior zebrafish neural tube. First, Myosin-driven apical constriction of midline neuroectoderm cells creates a medial hinge point and elevates the bilateral neural folds, producing a shallow groove along the dorsal midline. The neural folds come together near the center of this groove, pinching it into anterior and posterior segments. The neural folds zipper together posteriorly from the pinch point while the neural folds continue toward the midline in the anterior portion of the groove, creating the previously described eye-shaped opening that then zippers shut between two closure points. Around this time, the caudal-most end of the posterior portion completes its zipper closure at a distinct closure site (see model in **Fig. 7**).

**Figure 1.**
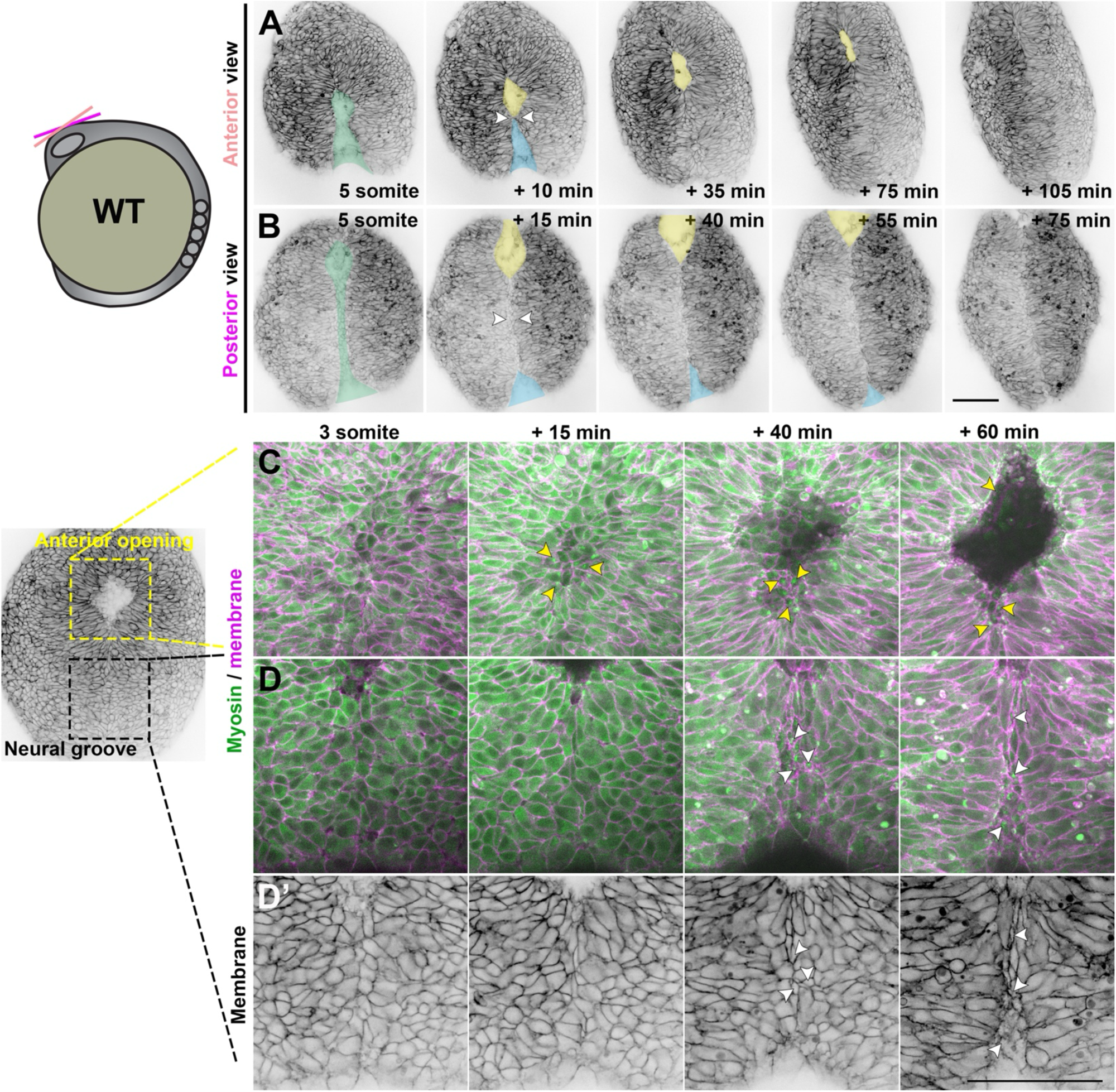
Neural fold fusion proceeds bidirectionally from a central “pinch point”. **A-B**) Still frames from time-lapse series of anterior neural tube development in WT or *tri* sibling embryos expressing membrane GFP or mCherry beginning at the 5 somite stage, viewed dorsally from more anterior (A) or posterior (B) positions. Green shading indicates the early neural groove, white arrowheads indicate the pinch point at which the bilateral neural folds make contact. Thereafter, yellow and blue shading indicate the anterior and posterior openings, respectively. Each image series is a single Z plane from a confocal stack and is representative of 30 individual WT and sibling embryos imaged in 8 independent trials. Additional examples are shown in Supp. Fig. 1 and Supp. videos 1-4. **C-D’**) Live images of the anterior (C) and posterior (D-D’) neural groove of a representative WT embryo expressing membrane Cherry (magenta in C-D, black in D’) and the Sf9-mNeon Myosin reporter (green) at the stages indicated. Arrows highlight Myosin localization to the medial apical cortex of apically constricted cells. The embryo image to the left is at the approximately + 50 minute time point. Anterior is up in all images, scale bars = 100 μm.

**Supplemental Figure 1.**
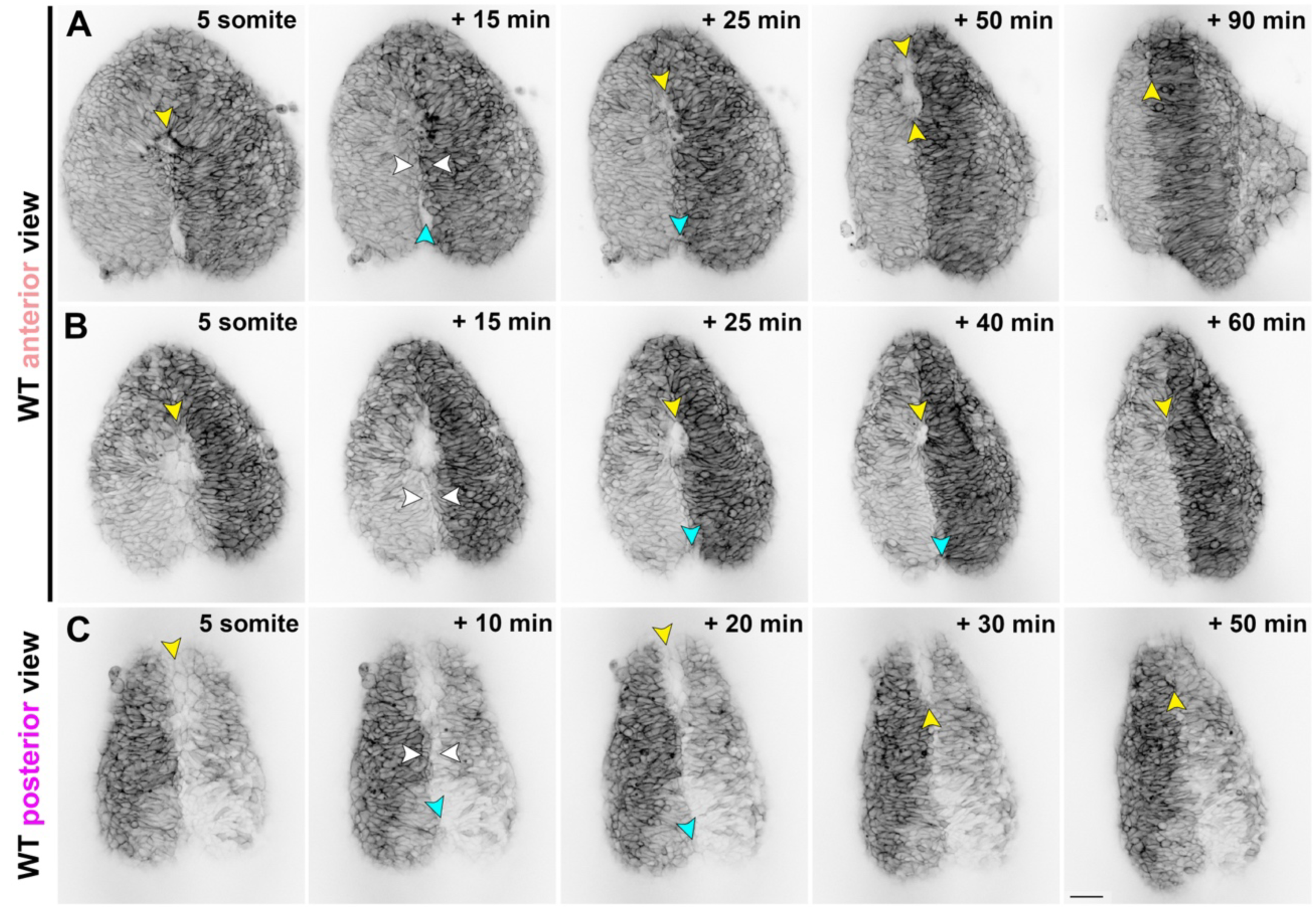
Live imaging reveals neural fold fusion dynamics in live WT embryos. (**A-C**) Still frames from time-lapse series of anterior neural tube development in WT or *tri* sibling embryos expressing membrane GFP or mCherry beginning at the 5 somite stage, viewed dorsally from more anterior (A-B) or posterior (C) positions. Yellow arrowheads indicate the anterior edge of the neural groove and eventually the eye-shaped opening. White arrowheads indicate the pinch point at which the bilateral neural folds make contact. Cyan arrowheads indicate the posterior opening that zippers closed in the posterior direction from the pinch point. Anterior is up in all images, scale bar = 50 μm.

### Neural fold fusion is abnormal in *vangl2* deficient embryos

Multiple aspects of primary neurulation, including convergent extension (CE) of the neural plate, apical constriction of the medial hinge point, and neural fold fusion, are regulated by planar cell polarity (PCP) signaling, and disruption of PCP signaling components prevents neural tube closure in multiple vertebrate models (*3, 40, 42, 43, 45, 46, 70–74*). Mutations in the core PCP gene *vangl2* also cause defects in CE and neurulation in zebrafish, but unlike the open neural tubes observed in other vertebrates, MZ*trilobite* (*tri*-/-) zebrafish embryos (lacking both maternally and zygotically expressed *vangl2*) instead present with ectopic bilateral neural lumens in the spinal cord. This phenotype is thought to result from delayed CE of the neural folds which causes C-divisions to occur laterally (*21, 22, 55*) rather than at the midline (*15, 17, 20*). To determine whether and how loss of *vangl2* disrupts neural tube closure in the forebrain region, we performed confocal time-lapse imaging of the anterior neural plate in zygotic *tri*-/-embryos. Because maternally expressed *vangl2* remains in *tri*-/-mutants, we also examined WT embryos injected with a translation-blocking morpholino oligonucleotide (MO) against *vangl2* (*75*), which also targets maternal *vangl2* transcripts (*27, 76*). Morphant phenotypes were largely rescued by *vangl2* mRNA lacking the MO binding site in the 5’ UTR (**Supp. Fig. 2**), highlighting its specificity.

We observed that the bilateral neural folds of both *tri*-/-mutant and *vangl2* morphant embryos began (at the 4 somite stage) much farther apart than heterozygous and WT siblings, which led to a delay in formation of the pinch point and ultimately the anterior eye-shaped opening. Indeed, while the neural groove in control embryos had pinched into anterior and posterior segments and formed the eye-shaped opening by 6-7 somite stage (**Fig. 1**), the neural folds of stage-matched *vangl2* mutants and morphants had often not yet made contact, leaving wide gaps between them (**Figs. 2-3**). The initial pinch point eventually formed around the 7-8 somite stage in many *vangl2* morphant and *tri-/-* embryos (approximately one hour delayed compared with controls), creating anterior and posterior openings (**Fig. 2A-C, Supp. videos 6-8**). As in WT, this pinching involved apical constriction of midline cells with Myosin localization at the medial apical cortex and edges of neural openings (**Fig. 2D, Supp. video 7**).

Unlike WT embryos, however, these openings did not always zipper smoothly closed. Instead, the neural folds in many *vangl2* deficient embryos were seen “buttoning up” at multiple discrete pinch points that formed in quick succession in the posterior (and in at least one example, the anterior) opening (**Fig. 2A-C**, white arrows, **Supp. Videos 6-7**). The result was a series of progressively more posterior openings that were themselves pinched in two by new “buttons”, which then closed by zippering bidirectionally away from the new pinch point (**Fig. 2A-C**, blue, indigo, pink shading). This was quantified as an increase in the number of both openings and pinch points observed in the neural tubes of *tri*-/- and (to a lesser extent) *vangl2* morphant embryos (**Fig. 2E-F**). Whether the additional pinch points are absent from WT embryos, or simply not visible in our live imaging conditions, is unclear. The neural tube closure defects in three *vangl2* morphant embryos were so severe that no pinch point formed during the imaging period (**Fig. 2F**), leaving a single large opening (**Fig. 2E**). Even in *vangl2* deficient embryos with pinch points, closure of the posterior-most opening was substantially delayed, yielding persistent openings visible in the posterior region of essentially all *vangl2* morphants and *tri*-/-mutants examined (**Fig 2A-C**, indigo and pink shading). Finally, rounded cells were seen protruding from the neural groove of most *vangl2* deficient embryos (**Fig. 2A, C, Fig. 3B, E**, orange arrows), which sometimes detached from the neural tube after closure (**Fig. 2C**, orange arrows). Together, these results highlight severe and regionally distinct defects in neural fold fusion in the absence of *vangl2*.

**Supplemental Figure 2.**
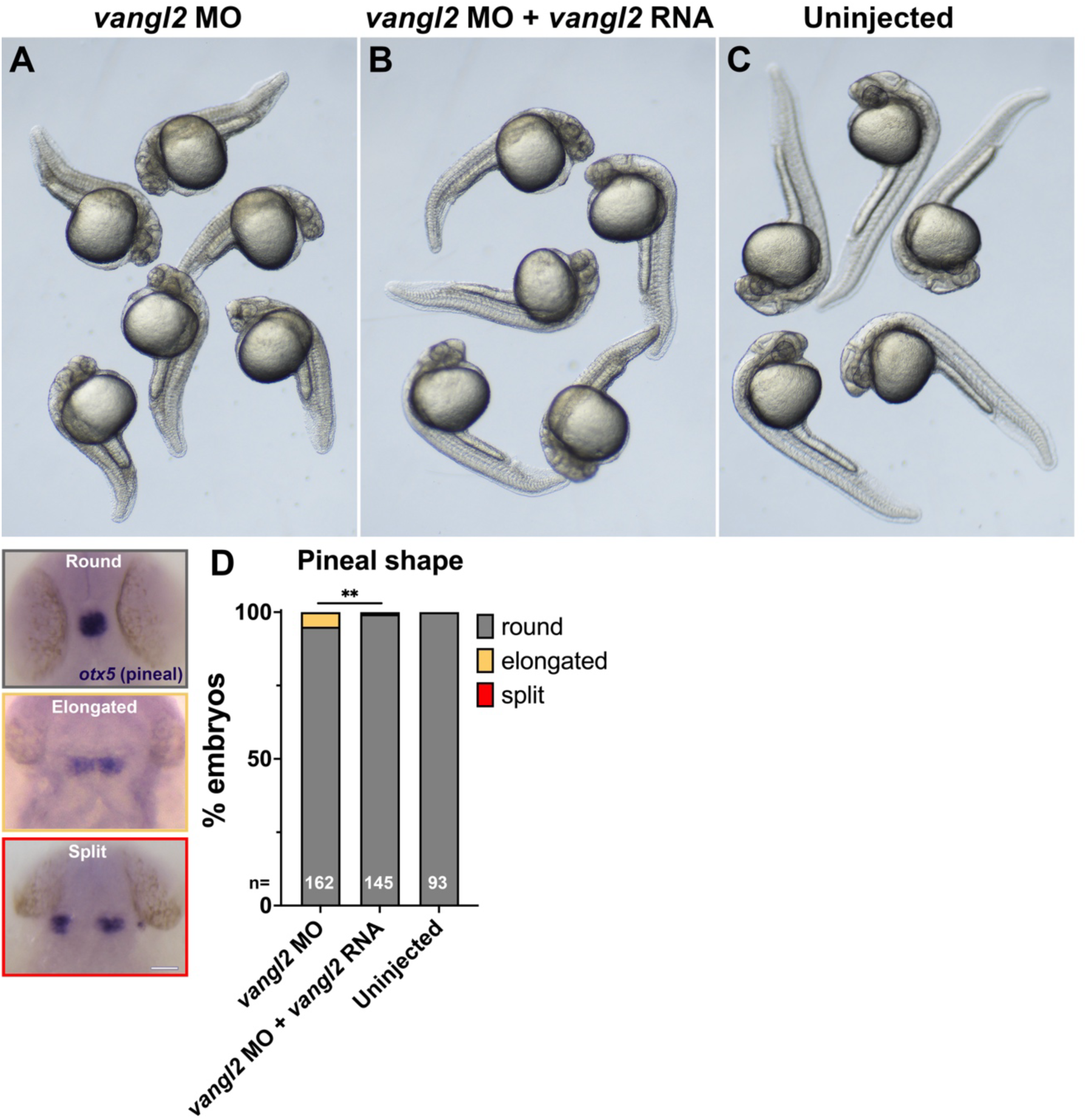
*vangl2* mRNA largely rescues *vangl2* morpholino day 1 phenotypes. (**A-C**) Live embryos at approximately 28 hpf injected at the 1-cell stage with 2 ng *vangl2* morpholino (A), 2 ng *vangl2* morpholino + 10 pg *vangl2* mRNA lacking the MO-binding site (B), or uninjected siblings. Images are representative of 3 independent trials. **D**) Classification of pineal shape in 28 hpf control, *vangl2* morphant, and rescued morphant embryos WISH stained for *otx5*. n values indicate the number of embryos of each condition measured from 3 independent trials. **p=0.0015, Fisher’s exact test.

**Figure 2.**
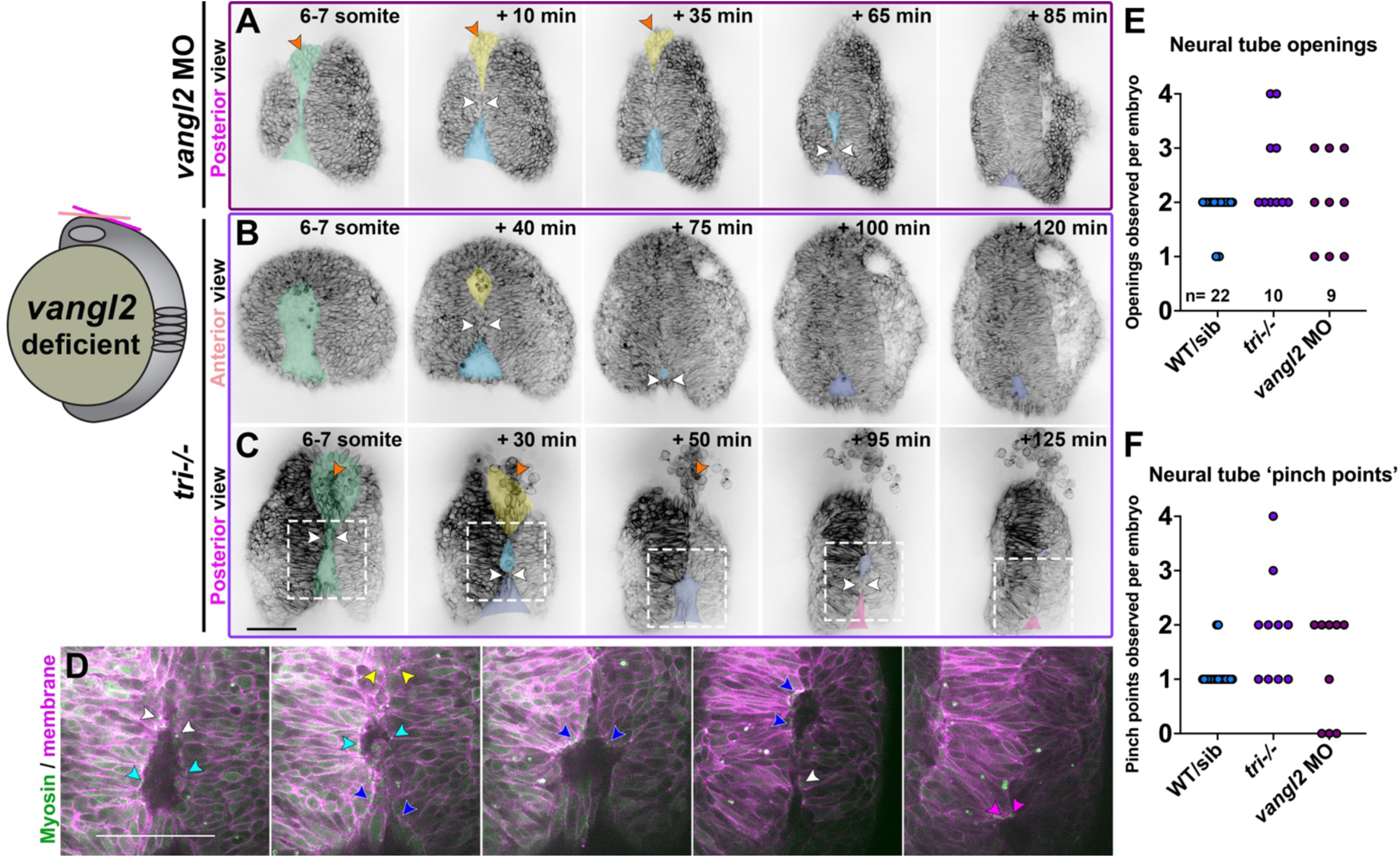
The neural folds of *vangl2* deficient embryos exhibit ectopic closure points. **A-C**) Still frames from time-lapse series of anterior neural tube development in *vangl2* morphant (A) or *tri-/-* embryos (B-C) expressing membrane GFP or mCherry beginning at the 5-somite stage, viewed dorsally from more anterior (B) or posterior (A, C) positions. Green shading indicates the early neural groove, white arrowheads indicate pinch points at which the bilateral neural folds make contact. Yellow and blue shading indicate the initial anterior and posterior openings, respectively. Indigo and pink shading indicate new openings formed by “pinching” of the initial posterior opening. Each image series is a single Z plane from a confocal stack and is representative of 8 morphant and 10 mutant embryos imaged in 2 or 4 independent trials, respectively. **D**) Enlargements of regions in (C) within dashed boxes showing membrane Cherry in magenta and the Sf9-mNeon Myosin reporter in green. Arrows highlight Myosin localization to the edges of neural fold openings. The colors of the arrowheads correspond to the color of shading in (C). **E-F**) Quantification of the number of neural fold openings (E) and pinch points (F) observed in time-lapse series of embryos of the conditions indicated. Each dot represents a single embryo from 2 WT, 2 *vangl2* morphant, and 4 *tri* independent trials. Anterior is up in all images, scale bars = 100 μm. See also Supp. videos 6-8.

### Anterior neural fold fusion is delayed in *vangl2* deficient embryos

Although a secondary pinch point was observed in the anterior neural opening of at least one *tri-/-* embryo, the anterior neural plate generally gave rise to a single large opening that ultimately zippered closed. Interestingly, while the eye-shaped opening of control embryos closed predominantly from anterior to posterior (**Supp. video 5**), the equivalent opening in *tri*-/-mutants and *vangl2* morphants often zippered closed from posterior to anterior (**Supp. video 8**). In both WT and *tri-/-* mutant embryos, the Sf9 Myosin reporter localized to the edges of the anterior eye-shaped opening (**Fig. 3A-B’**). Although its localization was dynamic, it concentrated near sites of zippering and around the circumference of the opening like a purse string (**Fig. 3A-B’**, yellow arrows), as observed during ascidian neural tube closure (*65*). Myosin remained localized at the closure site until the neural folds had fused, at which point it dissipated (**Supp. Video 4**).

As mentioned above, formation of the initial pinch point and the resulting anterior opening were substantially delayed in *vangl2* deficient embryos compared with WT and sibling control embryos, as was the subsequent closure of the anterior opening. This is reflected in quantitative measurements of the distance between the anterior neural folds over time, beginning at the 6 somite stage when the eye-shaped opening had formed in control embryos (**Fig. 3C-E**). A simple linear regression revealed that the distance between the neural folds of both *vangl2* morphants and *tri*-/-mutants started approximately two times larger (Y intercepts of 167.6 and 111.0 μm, respectively) than sibling control embryos (Y intercepts of 53.5 and 58.9) (**Fig. 3F**).

Interestingly, the rate of neural fold convergence was higher in *vangl2* morphants and mutants (with slopes of -0.76 and -0.48, respectively) compared with their sibling controls (slopes of -0.30 and -0.35) (**Fig. 3F**). This accelerated closure could not fully compensate for the increased width of their neural folds, however, and closure of the anterior opening was significantly delayed in *vangl2* morphants and mutants (with X intercepts at 220.7 and 231.3 minutes, respectively) with respect to sibling controls (X intercepts of 179.6 and 168.2 minutes) (**Fig. 3F**). This suggests that the zippering process itself is not disrupted in *vangl2* deficient embryos, and that delayed neural fold fusion may instead be the consequence of reduced CE of the neural plate (*21, 27, 55*).

**Figure 3.**
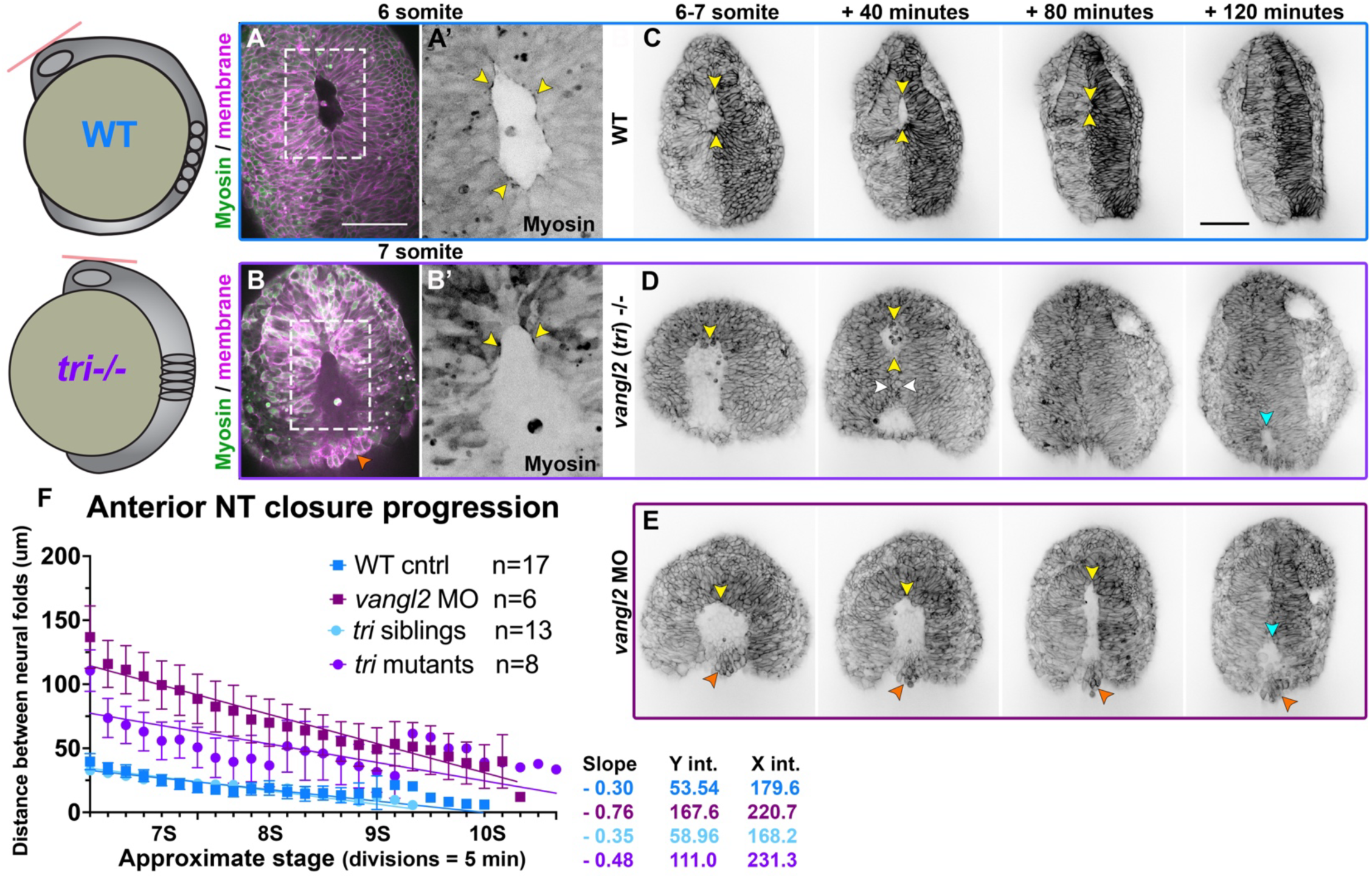
Neural fold fusion is delayed in *vangl2* deficient embryos. **A-B ’**) Live images of the anterior neural tube of WT (A) and *tri-/-* (B) embryos expressing membrane Cherry (magenta) and the Sf9-mNeon Myosin reporter (green in A-B, black in prime panels) at the stages indicated. Areas in dashed boxed are enlarged in the prime panels depicting Sf9-mNeon only. Yellow arrows highlight Myosin localization to the edges of the anterior neural opening. Orange arrow shows cells protruding from the opening of *tri-/-* embryos. **C-E**) Still frames from time-lapse series of anterior neural tube development in WT (C), *tri*-/-mutant (D), and *vangl2* morphant (E) embryos expressing membrane GFP or mCherry beginning at 6-7 somite stage, viewed dorsally. Yellow arrowheads indicate the anterior edge of the eye-shaped opening, white arrowheads indicate the pinch point, cyan arrowheads indicate the posterior opening, and orange arrowheads show cells protruding from the neural groove of *vangl2* morphants. Each image series is a single Z plane from a confocal stack and is representative of multiple embryos of that condition (see n values for each condition in F). **F**) Distance between the bilateral neural folds over time in embryos of the conditions indicated, beginning when the eye-shaped opening forms at the 6-somite stage. Symbols are mean + SEM, lines are simple linear regressions, for which slopes and intercepts are provided. n values indicate the number of embryos measured of each condition from 2 independent *vangl2* MO and 4 independent *tri* mutant trials. Anterior is up in all images, scale bars = 100 μm. See also Supp. videos 5, 8.

### The forebrain neural folds fuse to enclose a lumen

Neural tube closure in zebrafish and amniote embryos involves not only CE and zippering, but also formation of medial and dorsolateral hinge points and elevation of the neural folds (*8, 9, 12, 29, 77*). Using time-lapse confocal microscopy to collect optical transverse sections through the developing forebrain region, we observed hinge point formation and neural fold elevation in WT embryos beginning at the 3-4 somite stage (**Fig. 4A-C’**). As previously described (*29*), the forebrain neural plate began largely flat across the apical surface but developed a prominent medial hinge point by the 5 somite stage (**Fig. 4A-A’**). In the anterior region of the forebrain, cells lining the V-shaped neural groove “sealed up” progressively from ventral to dorsal until the neural tube was closed and smooth across its outer surface (**Fig. 4A**). This is apparent from measurements of medial hinge point angle in control embryos, which became more acute as the neural folds elevated (also as reported in (*29*)) and then widened slightly just as the folds sealed up (**Fig. 4E**).

Optical sections through a more posterior region of the forebrain, however, showed the bilateral neural folds elevating around a larger U-shaped groove. We found the Sf9-mNeon Myosin reporter accumulated at the apical surface of midline neural plate cells (**Supp. Fig. 3A’**), as previously described during hinge point formation in both the zebrafish forebrain and spinal cord (*28, 29*). Critically, the neural folds of WT embryos then fused at the dorsal side to enclose a lumen (**Fig. 4A’,** white arrow, **Supp. video 9**) in a fashion strikingly similar to primary neurulation in amniote embryos. Myosin accumulated along the apical surface of the deepening neural groove and remained until the neural folds fused (**Supp. Fig. 3A’**). This is distinct from the mechanism of lumen formation in the zebrafish spinal cord, which occurs through cavitation of the solid neural rod (*16, 23*). We also observed that at this more posterior position, the periderm separated slightly from the underlying ectoderm and bridged the gap between the bilateral neural folds until they fused dorsally (**Fig. 4A’**, orange arrow), which can also be observed in time-lapse series from a previous study (*29*). These data directly demonstrate that neural folds within the forebrain region of zebrafish embryos elevate and fuse to enclose a lumen, highlighting conservation of fold-and-fuse neurulation across vertebrates.

### Neural groove formation and closure is abnormal in *vangl2* deficient embryos

Our live confocal imaging revealed significant delays in neural fold fusion in *vangl2* deficient embryos (**Fig. 3**), but it was unclear whether this delay alone underlies the large openings we observed in their forebrain regions. To this end, we collected transverse optical sections through the developing brains of *tri*-/-mutant and *vangl2* morphant embryos. The anterior forebrain regions of *tri*-/-embryos (**Fig. 4B**) were wider than their siblings throughout neural tube development (**Fig. 4D**) but exhibited formation of a V-shaped groove that “sealed up” from ventral to dorsal, similar to (but forming a less acute angle) than sibling controls (**Fig. 4E**). The posterior forebrain regions of *tri*-/-embryos also exhibited U-shaped grooves, but these were substantially larger by cross-sectional area than sibling controls (**Fig. 4F**), likely reflecting increased width of the neural plate. Unlike WT embryos, however, this region of the neural tube did not fuse to enclose a lumen, but instead “sealed up” from ventral to dorsal as in the more anterior regions of both *tri-/-* mutants and siblings (**Fig. 4B’, Supp. Video 10**). Despite increased width of the neural groove and its abnormal method of closure, Myosin localization appeared largely normal within *tri*-/-embryos, accumulating at the apical surface of cells within the deepening groove (**Supp. Fig. 3B’**).

*vangl2* morphants presented with a more severe phenotype than *tri*-/-mutants. The anterior forebrain regions of morphants were even wider than mutants (**Fig. 4D**) and exhibited neither hinge points nor V-shaped grooves, instead resembling a solid mass of cells at this level (**Fig. 4C, E**). A large U-shaped groove was apparent in more posterior regions, but fusion of the neural folds was significantly delayed and sometimes blocked entirely (**Fig. 4C’, Supp. video 11**), as evidenced by the enlarged cross-sectional area of the neural groove over time (**Fig. 4F**). Notably, cross-sectional area of the neural groove was larger in *vangl2* mutants than morphants at early stages (**Fig. 4F**), likely reflecting reduced hinge point formation in morphants. In both *tri*-/-mutants and *vangl2* morphants, the periderm spanned the gap between neural folds in the posterior region as in control embryos, although this cell layer separated from the underlying neural plate earlier and by a larger distance than in controls. These cells were also rounded and protruded outward from the neural groove (**Fig. 4B’-C’**, orange arrows), indicating that cells observed protruding from the neural groove of *vangl2* deficient embryos (**Figs. 2 and 3**) are periderm that adopted an abnormal shape. Together, these results suggest that increased width of the neural plate, and not absence of Myosin localization, is likely primarily responsible for larger neural grooves in the forebrain region of *vangl2* deficient embryos. The neural folds surrounding these enlarged grooves are both delayed in their closure and utilize a different fusion method than WT embryos, sealing up from ventral to dorsal rather than enclosing a hollow lumen.

**Figure 4.**
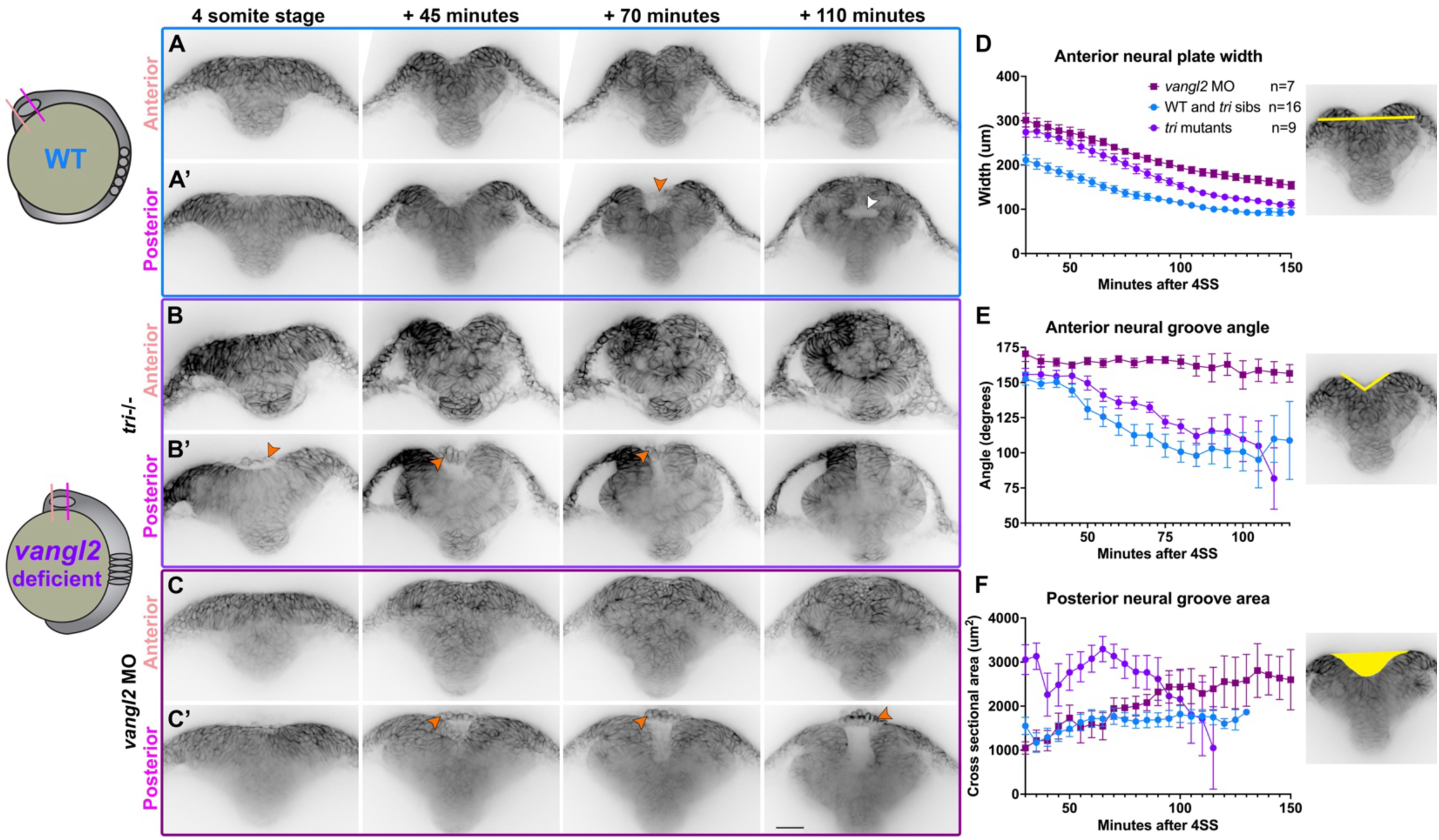
The bilateral neural folds fuse dorsally to enclose a lumen in WT, but not *vangl2* deficient embryos. (**A-C ’**) Still frames from time-lapse series of neural fold fusion in WT (A), *tri*-/-mutant (B), and *vangl2* morphant (C) embryos expressing membrane GFP or mCherry beginning at the 4-somite stage, viewed in transverse optical section through anterior (A-C) or posterior (A’-C’) regions of the forebrain. Orange arrowheads indicate periderm cells spanning the neural groove. Each image series is a single representative Z plane from a confocal stack. **D-F**) Measurements of neural plate width (D) and neural groove angle (E) within the anterior forebrain region and cross-sectional area of the neural groove (F) within the posterior forebrain region in embryos of the conditions indicated, beginning at 4-somite stage. Symbols are mean + SEM, n values indicate the number of embryos measured of each condition from 2 independent *vangl2* MO and 3 independent *tri* mutant trials. Control embryos for MO and mutant experiments were combined in graphs (blue lines) for simplicity. Images to the right are illustrative of the measurements made. Dorsal is up in all images, scale bar = 50 μm. See also Supp. videos 9-11.

**Supplemental Figure 3.**
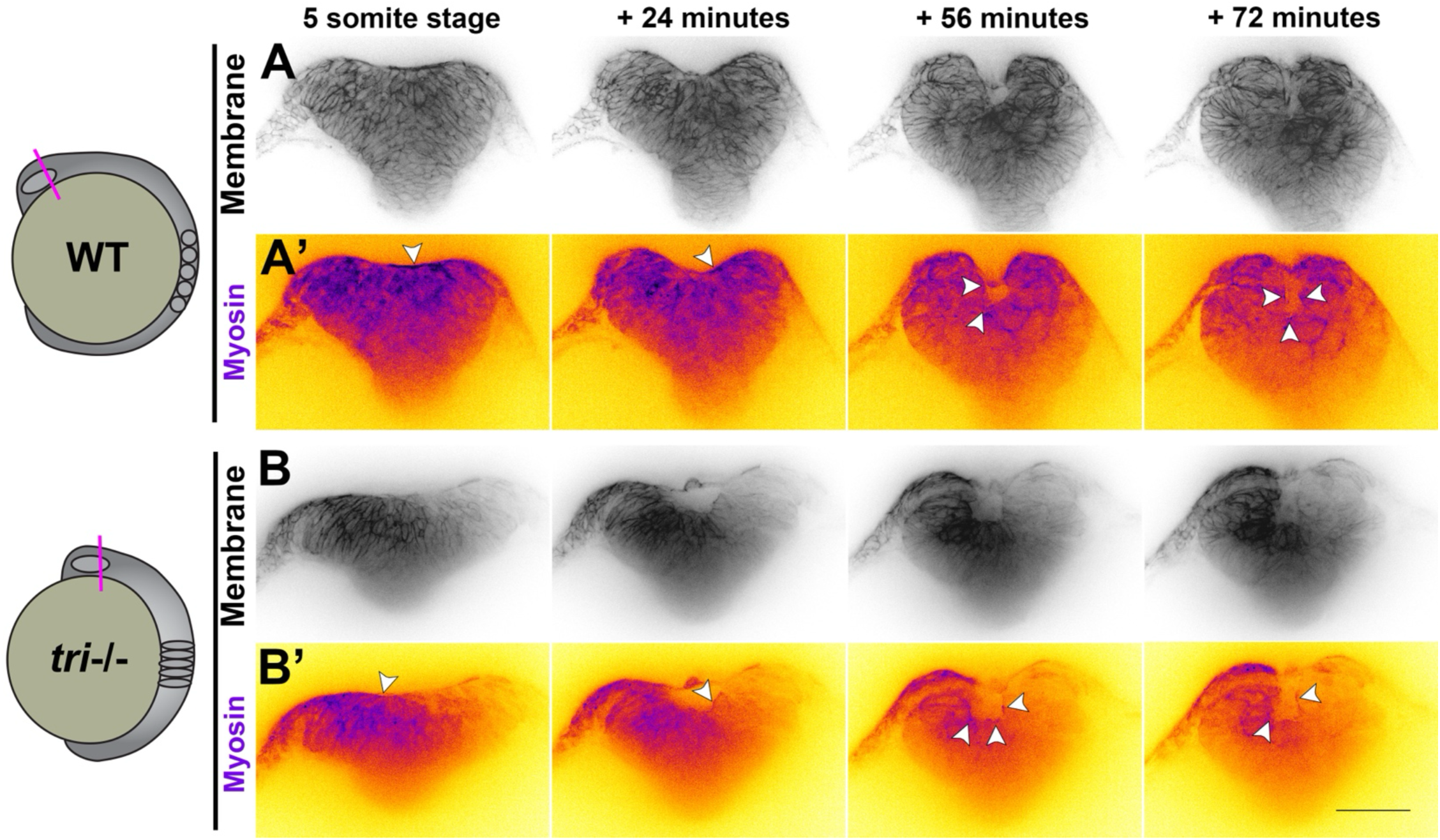
Myosin accumulates apically within the neural groove of WT and *vangl2/ tri-/-* mutant embryos. (**A-B’**) Still frames from time-lapse series of neural fold fusion in WT (A) and *tri*-/-mutant (B) embryos expressing membrane mCherry (A-B, black) and Sf9-mNeon (A’-B’, inverted Fire look up table) beginning at the 5-somite stage, viewed in transverse optical section through the posterior region of the forebrain. Arrowheads indicate Myosin localization to the apical surfaces of cells comprising the neural groove. Each image series is a single Z plane from a confocal stack representative of 4 sibling and 4 mutant Sf9-expressing embryos. Dorsal is up in all images, scale bar = 100 μm.

### Day-one phenotypes poorly reflect abnormal neural tube closure in *vangl2* deficient zebrafish

Our identification of fold-and-fuse neurulation within the zebrafish forebrain region, combined with defects in this process upon loss of the *vangl2*, a homolog of an NTD risk gene (*53, 54*)), raises the possibility of modeling NTDs in zebrafish. However, live imaging is too time- and labor-intensive for rapid screening of multiple risk genes, an approach to which zebrafish is otherwise well suited (*78, 79*). We therefore examined fixed *vangl2* deficient embryos for hallmarks of defective or delayed neural tube closure. Bifurcation of pineal gland precursors and the dorsal roof plate around 24 hours post fertilization (hpf) were previously suggested as proxies for NTDs in zebrafish embryos with disrupted Nodal signaling or mutations in *cdh2* (encoding N-cadherin) (*30–33*). To determine whether similar phenotypes were present in zebrafish embryos lacking *vangl2*, we examined the morphology of these structures in *tri*-/-mutants, *vangl2* morphants, and sibling controls at approximately 28 hpf. Both a transgenic *flh:kaede* (*80*) line (**Fig. 5A**) and whole mount *in situ* hybridization (WISH) for the pineal marker *otx5* (**Fig. 5B**) revealed a small subset of *vangl2* morphants with elongated or split pineal domains which were never observed in control siblings (**Fig. 5C**). These phenotypes were significantly rescued by co-injection of morpholino-resistant *vangl2* mRNA (**Supp. Fig. 2D**), demonstrating they are caused by loss of *vangl2* specifically. Notably, *tri*-/-embryos did not exhibit abnormal pineal morphology at this stage (**Fig. 5D**), which is likely attributable to maternally deposited *vangl2* that is eliminated by the MO (*27, 76*). WISH for the dorsal neural tube marker *wnt1* also revealed bifurcated roof plates in a subset *vangl2* morphants (**Fig. 5E-H**). Histological analysis of 28 hpf *vangl2* morphant neural tubes (fore-through hind-brain) with split or elongated pineal domains revealed ectopic tissue and supernumerary midline structures (**Fig. 5I-L**, yellow arrows) similar to those reported in the spinal cord region of *vangl2* deficient embryos (*21, 22, 55*). All fifteen embryos examined contained ectopic midline tissue caudally (**Fig. 5J-L**) and nine exhibited this defect at the level of the pineal (**Fig. 5J-K**), suggesting that split pineal domains are associated with ectopic midline tissues resulting from abnormal C-divisions.

We compared these phenotypes to those of Nodal signaling-deficient embryos, in which split pineal domains were proposed to represent open neural tubes (*30–32*). As previously reported, embryos lacking Nodal signaling through maternal and zygotic loss of the coreceptor *tdgf1/oep* (MZ*oep*-/-) or treated with the Nodal inhibitor SB505124 exhibited substantially wider and split pineal domains (**Supp. Fig. 4A-E**). Histological analysis of SB505124-treated embryos at 28 hpf revealed a striking Swiss cheese-like pattern of multiple small holes in the neural tube of every embryo examined, regardless of whether their pineal domains were split (**Supp. Fig. 4F-H**). This internal anatomy of the neural tube is consistent with previous reports of multiple lumen-like structures in MZ*oep*-/-embryos (*81*), but is distinct from the ectopic bilateral midlines observed in *vangl2* morphants (**Fig. 5J-L**) despite similar pineal phenotypes. This demonstrates that bifurcation of the pineal precursors can be underlain by multiple distinct internal phenotypes, which do not appear to share features with amniote NTDs. Furthermore, although all *tri*-/-mutant and *vangl2* morphant embryos examined by live imaging exhibited defects and delays in neural tube closure, only a small fraction of morphant and no *tri*-/-mutant embryos had split or elongated pineal domains at 28 hpf. Together, these results suggest that day-one pineal morphology is not representative of earlier neurulation phenotypes and therefore, not an ideal proxy for neural tube defects in zebrafish.

**Figure 5.**
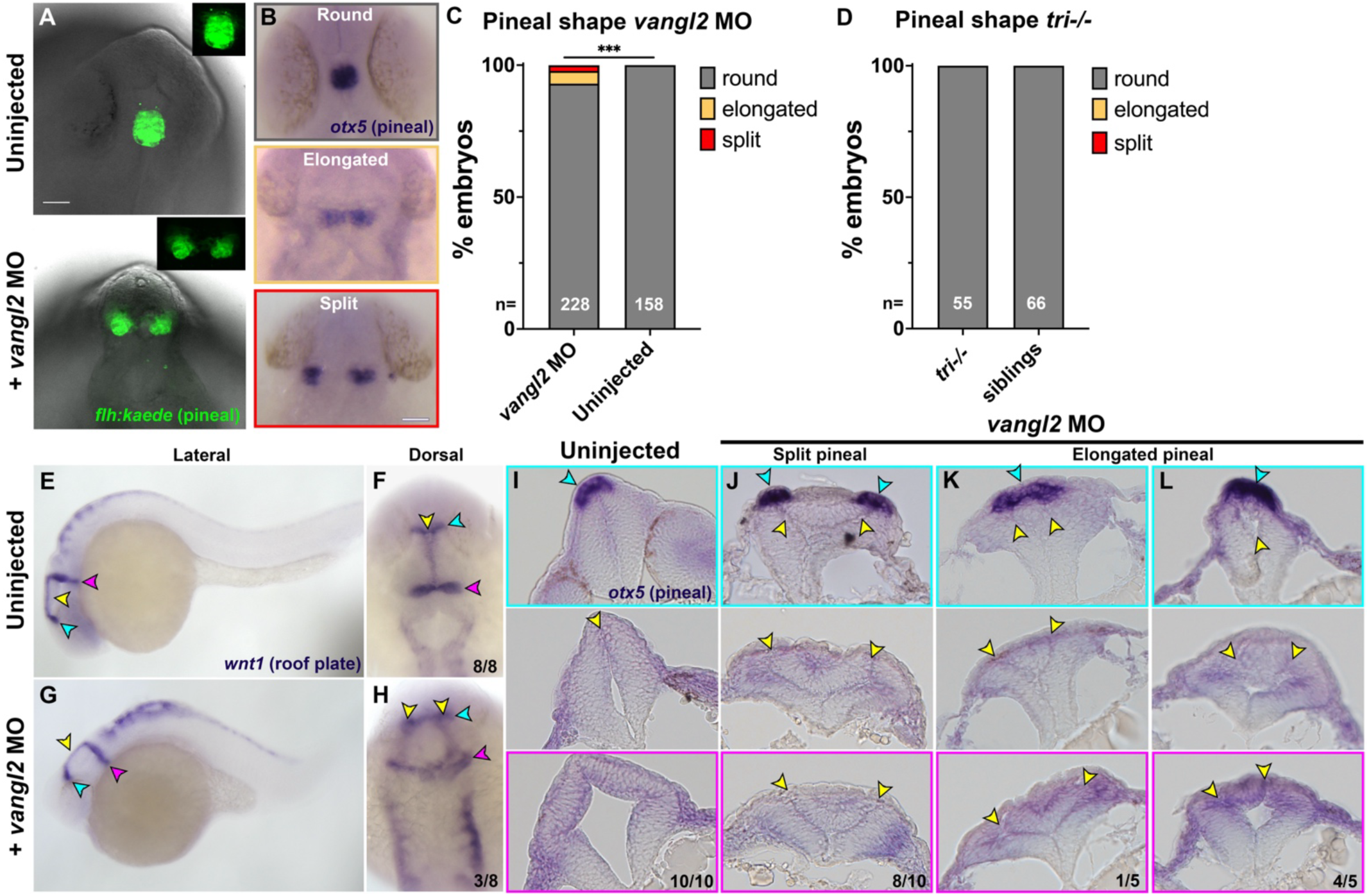
Pineal and roof plate morphology are disrupted in a subset of *vangl2* deficient embryos. (**A**) Example images of pineal precursor morphology in Tg[*flh:kaede*] control (top) or *vangl2* MO-injected (bottom) embryos at 28 hpf, viewed from the dorsal side. **B**) Examples of the three classes of pineal precursor morphology visualized by WISH for *otx5*. **C-D**) Classification of pineal shape in 28 hpf control and *vangl2* MO-injected (C) or *tri*-/-mutant (D) embryos expressing *flh:kaede* or WISH stained for *otx5*. n values indicate the number of embryos of each condition measured from 3 independent trials. ***p=0.0006, Fisher’s exact test. Scale bars = 50 μm. **E-H**) WISH for roof plate marker *wnt1* at 28 hpf in control (E-F) and *vangl2* MO-injected (**G-H**) embryos. Yellow arrowheads indicate the midline roof plate, cyan arrowheads indicate the epithalamus, and magenta arrowheads indicate the mid-hindbrain boundary (MHB). **I-L**) Transverse histological sections through the anterior neural tube at the level of the epithalamus (top panels), midbrain (middle panels), and MHB (bottom panels) in 28 hpf embryos of the conditions indicated. Cyan arrowheads indicate pineal precursors stained by *otx5* WISH. Yellow arrowheads indicate the neural tube midline(s)/lumen(s). Fractions indicate the number of embryos with the depicted phenotype over the total number of embryos examined for each condition. Anterior is up in (A-B, F, H), dorsal is up in (E, G, I-L).

**Supplemental Figure 4.**
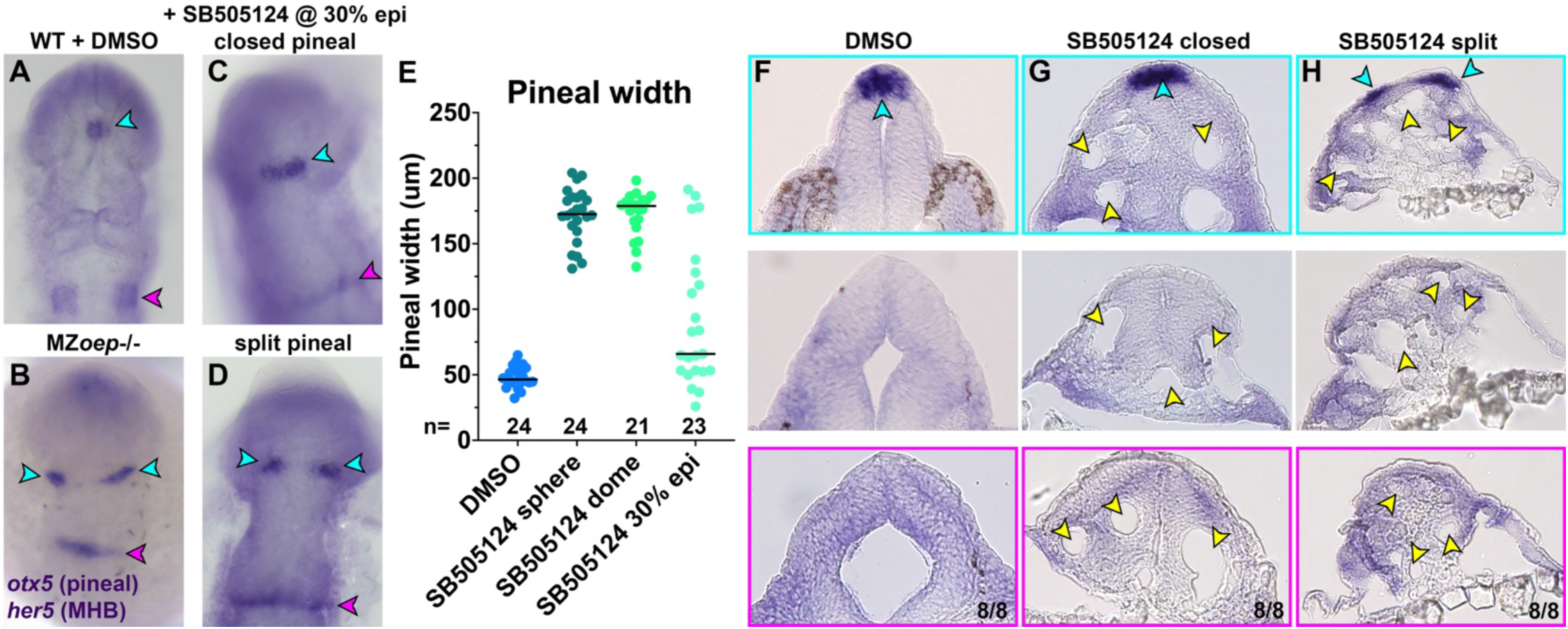
Distinct neural tube morphologies underlie split pineal phenotypes in Nodal deficient embryos. **A-D**) Representative images of the anterior neural tube in DMSO-treated WT (A), MZ*oep*-/-, or SB505124-treated (C-D) embryos at 28 hpf WISH stained for *otx5* and *her5*, viewed dorsally. Cyan arrowheads indicate pineal precursors, magenta arrowheads indicate the MHB. **E**) Width of pineal precursor domains in embryos of the conditions indicated, measured from *otx5* WISH at 28 hpf (as shown in A-D). Each dot represents a single embryo, black bars are median values. **F-H**) Transverse histological sections through the anterior neural tube at the level of the epithalamus (top panels), midbrain (middle panels), and MHB (bottom panels) in 28 hpf embryos of the conditions indicated. Cyan arrowheads indicate pineal precursors stained by *otx5* WISH. Yellow arrowheads indicate ectopic lumens in a Swiss cheese-like pattern. Fractions indicate the number of embryos with the depicted phenotype over the total number of embryos examined for each condition. Anterior is up in (A-D), dorsal is up in (F-H).

### Neural tube defects are apparent in fixed *vangl2* deficient embryos

Because the neural tube abnormalities observed by live imaging in *vangl2* deficient embryos were not reflected in pineal morphology of one-day-old embryos, we examined pineal precursors in fixed embryos at earlier (3-10 somite) stages. In WT siblings from *tri+/-* incrosses and uninjected WT controls, pineal precursors marked by *flh/noto* expression began as bilateral domains that converged at the midline and fused at approximately 7 somite stage (**Fig. 6A, D**), as previously described (*30*). This convergence was delayed in *tri-/-* mutants (and in some *tri*+/-heterozygotes), as reflected by significantly wider *flh*+ domains from 3-4 somite stages (**Fig. 6B, D**). Pineal precursor widths trended wider in *tri* mutants and heterozygotes until the 8 somite stage, but were no longer statistically significant. The pineal precursors of *vangl2* morphants, however, were significantly wider than uninjected controls at all stages examined (**Fig. 6C, E**), consistent with more severe phenotypes in *vangl2* morphants than mutants (**Figs. 3-5**). We again speculate that this is due to blocked translation of maternal *vangl2* transcripts, which is supported by our finding that the severity of pineal phenotypes increases with increasing doses of *vangl2* MO (**Supp. Fig. 5**).

In addition to more widely-spaced pineal precursors, we observed pit-shaped openings in the anterior neural plate between of fixed *tri*-/-mutant and *vangl2* morphant embryos (**Fig. 6B-C, F-G**). These adopted a variety of shapes that we assigned to each of 5 categories increasing in severity from no apparent opening or 3D structure (category 0) to a deep groove that extended posteriorly from the pineal domain (category 4). The majority of WT embryos examined belonged to category 0, although we observed categories 1-2 (and occasionally 3) at the earlier stages examined (**Fig. 6F-G**), likely reflecting the neural grooves and openings found in all embryos prior to neural fold fusion. Category 4 neural tubes, however, were only observed in *tri*-/-mutant and (more commonly) in *vangl2* morphant embryos (**Fig. 6F-G**). Their incidence decreased as development proceeded, with most *vangl2* deficient embryos exhibiting category 0 phenotypes by 8 somite stage (**Fig. 6F-G**), likely reflecting the (delayed) neural fold fusion we observed by live imaging. 3-dimensional reconstructions of the forebrain region of fixed and phalloindin-stained *tri*-/-mutant and *vangl2* morphant embryos (**Fig. 6H-J**) corroborated our WISH images. While the dorsal surface of WT and control embryos appeared smooth at all stages examined, approximately half of *tri*-/-mutants and all *vangl2* morphant embryos examined exhibited pit-like structures (**Fig. 6I-J**, yellow arrowheads) from which rounded periderm cells protruded, as seen in our live imaging. Many *vangl2* deficient embryos also possessed a second, more posteriorly positioned pit structure (**Fig. 6I-J**, cyan arrowheads). Together, these data indicate that the widened neural grooves and delayed neural fold fusion we observed in live *vangl2* deficient embryos are also visible in fixed embryos, providing a simple readout for anterior neural tube defects in zebrafish.

**Figure 6.**
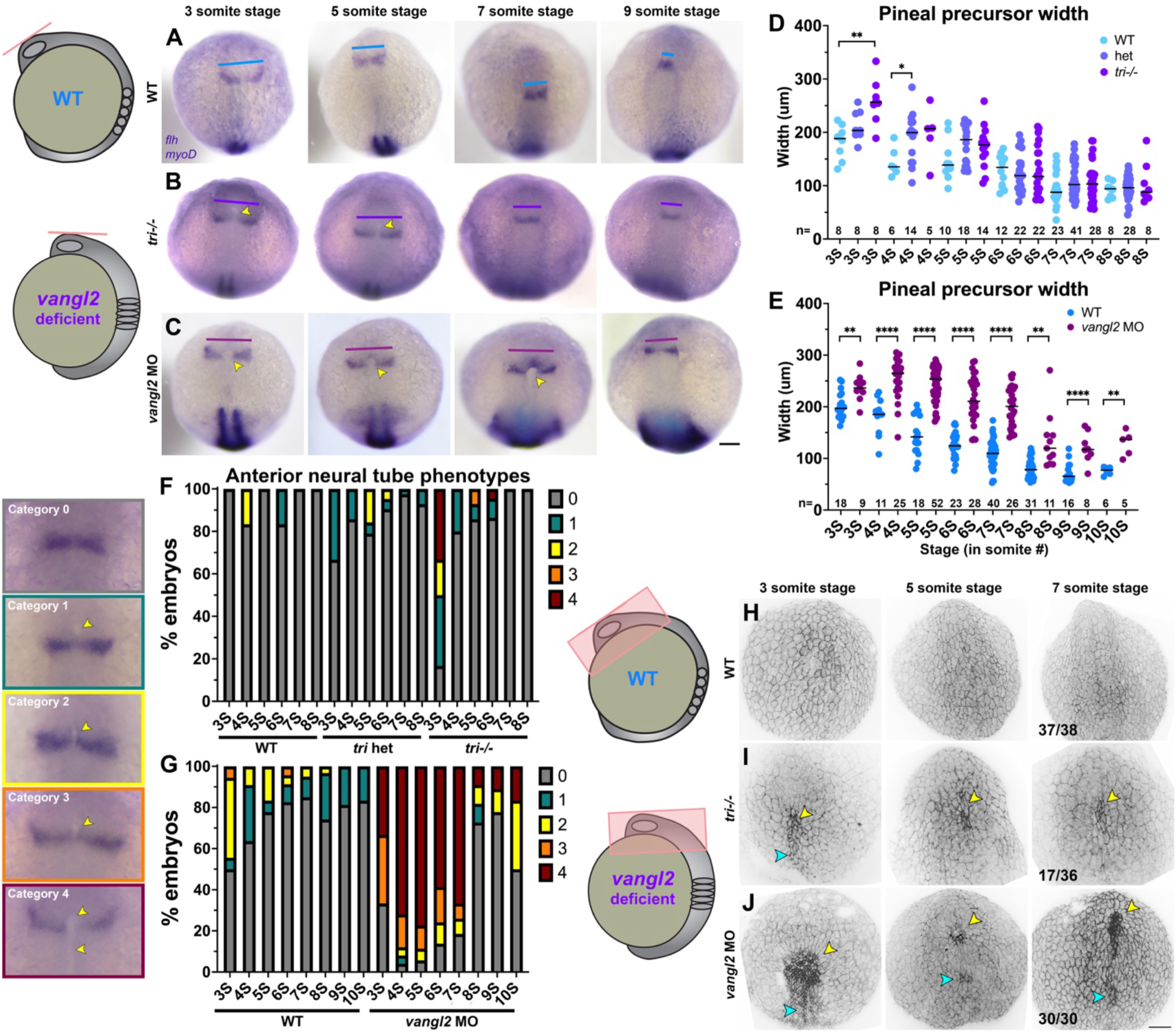
Fixed *vangl2* deficient embryos exhibit apparent delayed pineal convergence and openings in the anterior neural plate. (**A-C**) Representative images of pineal precursors (*flh* WISH) and somites/adaxial cells (*myoD* WISH) in control (A), *tri-*/-mutant (B), and *vangl2* MO-injected (C) embryos at the stages indicated, viewed dorsally. Blue, purple, and burgundy lines indicate pineal precursor width, yellow arrowheads indicate apparent openings in the anterior neural plate. **D-E**) Width of pineal precursor domains (as shown in A-C) in embryos from *tri+/-* incrosses (D) and in control (blue) and *vangl2* morphant (burgundy) embryos (E) at the stages indicated. Each dot represents a single embryo, black bars are median values. n values indicate the number of embryos of each stage/condition measured from 3 independent trials, **p=0.005, *p=0.045, Welch’s ANOVA with Dunnett’s multiple comparisons (D), **p<0.01, ****p<0.0001, multiple T-tests (E). **F-G**) Percentage of *tri*+/-incross (F) or *vangl2* morphant and control (G) embryos (as shown in A-C) at the stages indicated exhibiting the categories of anterior neural plate phenotypes shown on the left. n values as in D-E. **H-J**) 3-dimensional reconstructions of confocal Z-stacks through the anterior neural plate of fixed and phalloidin-stained embryos, viewed dorsally, of the stages and conditions indicated. Yellow arrowheads indicate apparent anterior openings, cyan arrowheads indicate openings in more posterior regions. Fractions indicate the number of embryos with the pictured phenotype over the number of embryos examined for each condition from 3 independent trials. Anterior is up in all images, scale bars = 100 μm.

**Supplemental Figure 5.**
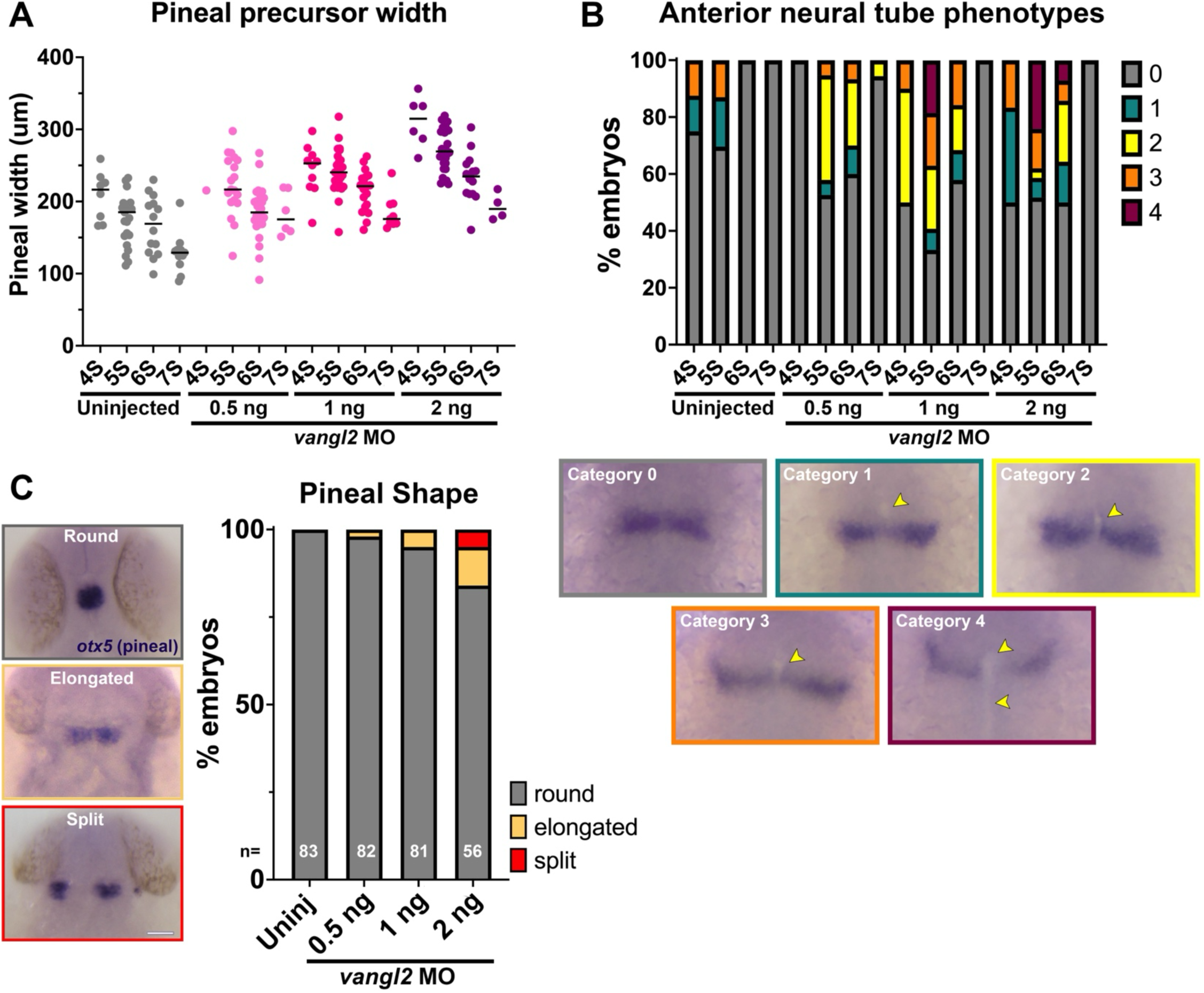
Severity of neural tube phenotypes increases with dose of *vangl2* MO. **A)** Width of pineal precursor domains in embryos injected with increasing doses of *vangl2* morpholino and uninjected controls at the stages indicated. Each dot represents a single embryo from 3 independent trials, black bars are median values. **B**) Percentage of embryos injected with each dose of *vangl2* morpholino at the stages indicated exhibiting the categories of anterior neural tube phenotypes shown below. **C**) Classification of pineal shape in 28 hpf control and *vangl2* MO-injected embryos WISH stained for *otx5*. n values indicate the number of embryos of each condition measured from 3 independent trials. ***p=0.0004, Fisher’s exact test. Anterior is up in all images.

### Anterior neural tube openings correlate with severity of convergent extension defects

*Vangl2* deficient embryos have well described defects in CE morphogenesis during gastrulation (*27, 61, 76*), and their neural tube phenotypes at later stages are thought to be secondary to reduced CE (*21*). Because our analysis of live and fixed *vangl2* deficient embryos revealed widened neural plates at the time of neural tube closure, we hypothesized that defective CE also underlies the observed delay in convergence and fusion of the bilateral neural folds. Indeed, *myoD*+ somites were significantly wider (consistent with reduced CE) in *tri*-/-mutants and *vangl2* morphants than their WT or heterozygous siblings (**Supp. Fig. 6A-B**), as described previously (*61, 76, 82, 83*). We noted that *vangl2* morphants, more of which exhibited severe neural tube phenotypes than *tri*-/-mutants, also tended to have wider somites than *tri*-/-mutants (**Supp. Fig. 6A-B, G**). And although not statistically significant, *vangl2* morphant embryos with more severe category 3-4 neural tube phenotypes trended toward wider somites than morphants with less severe category 0-2 phenotypes (**Supp.** Fig. 6C-G). These findings suggest that embryos with more severe CE defects are more likely to exhibit abnormal openings in the neural tube, which combined with our observations from live imaging, implicate reduced CE in delayed/abnormal neural tube closure upon loss of *vangl2*. We further speculate that some *vangl2* morphants can never overcome this delay to fuse the anterior neural folds before the onset of C-divisions around 10-somite stage, which go on to become the minority of embryos with split pineal precursors and/or roof plates at 28 hpf (**Fig. 5**).

**Supplemental Figure 6.**
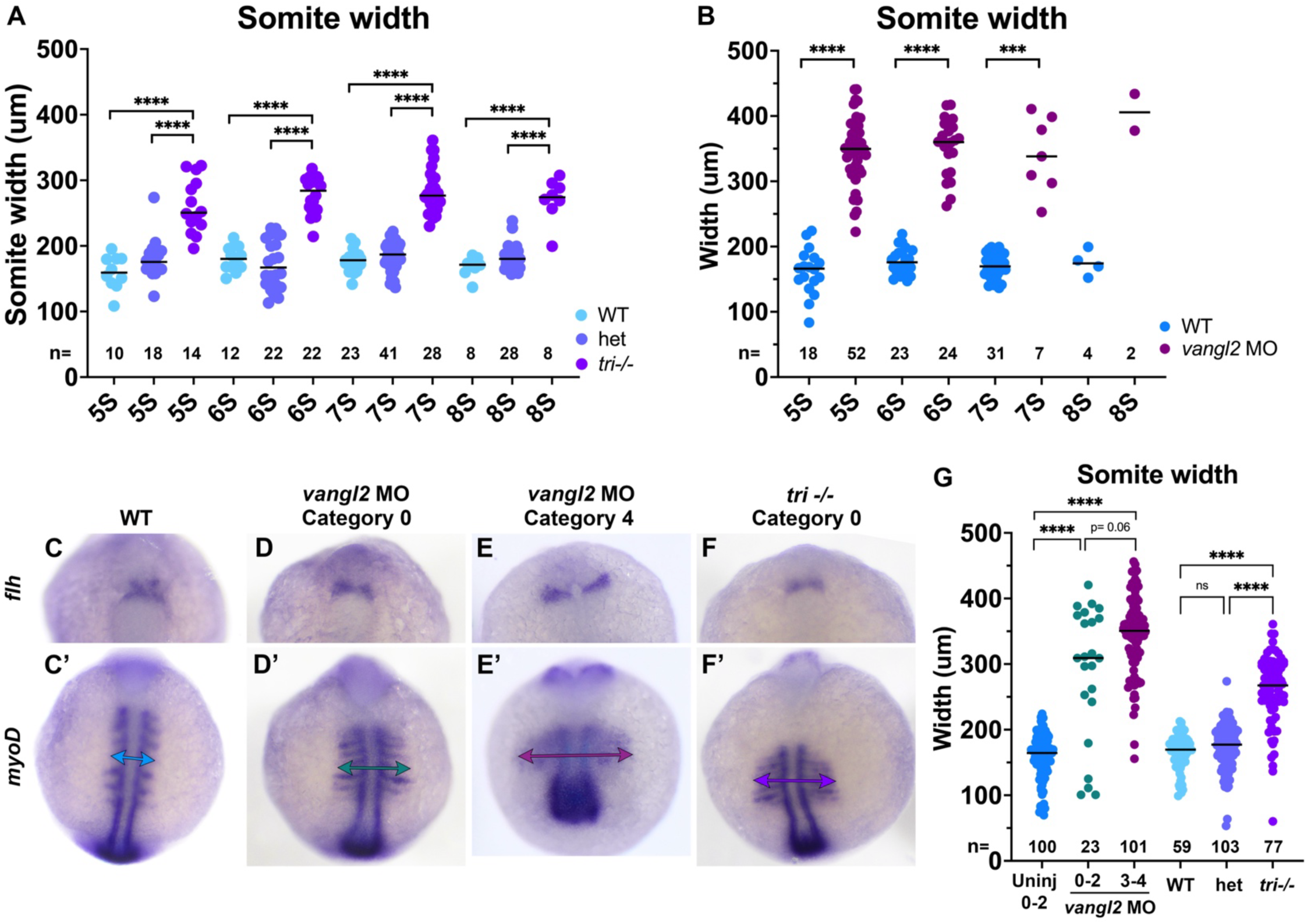
Anterior neural plate phenotypes are correlated with somite width. **A-B**) Width of *myoD*+ somites in *tri*+/-incross (A) and *vangl2* morphant and control (B) embryos at the stages indicated. ****p<0.0001, Welch’s ANOVA with Dunnett’s multiple comparisons (A), ****p<0.0001, ***p=0.0002, multiple T-tests with Welch’s correction (B). **C-F’**) Representative images of pineal precursors (*flh* WISH) and somites/adaxial cells (*myoD* WISH) in embryos of the condition and with the category of neural plate phenotype indicated at the 7-somite stage. Double arrows indicate somite width. **G**) Somite width (as shown in C’-F’) in embryos of the conditions indicated at 3-7 somite stages, including *vangl2* morphant embryos with category 0-2 (teal) or category 3-4 (burgundy) neural tube phenotypes. Each dot represents a single embryo, black bars are median values. ****p<0.0001, Kruskal-Wallis test with Dunn’s multiple comparisons. n values indicate the number of embryos of each stage/condition measured from 3 independent trials. Anterior is up in all images.

## Discussion

Closure of the neural tube is essential for proper development of the central nervous system, and its failure leads to deadly and debilitating congenital anomalies. Primary neurulation is well described in vertebrate models including mouse, chick, and *Xenopus*, which share a core set of cellular behaviors including convergent extension of the neural plate, apical constriction at hinge points, and dorsal fusion of the bilateral neural folds. Although neurulation in zebrafish embryos differs outwardly from these other species, it is increasingly clear that many aspects of primary neurulation are conserved, including apical constriction of neural midline cells and zippering of the neural folds (*23, 28, 29*). In this study, we identified additional commonalities between neural tube development in zebrafish and other vertebrate species – namely fold-and-fuse neurulation – and characterized their disruption by loss of the NTD risk gene *vangl2*.

### Conservation of primary neurulation mechanisms

It has long been appreciated that the zebrafish trunk neural tube forms through in-folding, by which the lateral edges of the neural plate come together at the dorsal surface (*17, 84*). This is facilitated by an enrichment of myosin contractility and subsequent apical constriction of midline cells, which drives their internalization (*28*). Although these features are common to other vertebrate embryos, they differ in that the neural tube lumen forms in mice and chick when it is enclosed by the bilateral neural folds upon dorsal fusion, whereas the zebrafish lumen (at the level of the trunk) forms later by cavitation of a solid rod (*16*). Hypotheses for evolutionary drivers of this unique method of neurulation include reducing exposure of the neural tube lumen to the outside environment (*85*) and overcoming the high mechanical stress imposed by axial curvature of the embryo that could otherwise prevent elevation of the neural folds (*86*). However, our findings provide direct evidence for neural fold elevation and enclosure of a lumen in zebrafish. Our time-lapse microscopy directly demonstrates that, within the future forebrain, the bilateral neural folds elevate around a midline groove then fuse at the dorsal surface, producing a hollow lumen (**Fig. 4**, **Fig. 7**). This mechanism is apparently unique to a portion of the forebrain region, as it was not observed in transverse images through more anterior (**Fig. 4**) or posterior regions (*16, 28, 84, 87*) of the neural keel, which is likely why it was not described previously. An elegant live imaging study did capture formation of hinge points and elevation of neural folds in the forebrain region (*29*), but did not describe lumen enclosure.

Our findings also expand our understanding of neural fold fusion within zebrafish. The aforementioned live imaging study of zebrafish forebrain (*29*) directly demonstrated neural fold fusion by bidirectional “zippering” of an eye-shaped opening. By imaging an earlier stage of neural development, the current study captures the events preceding formation of this closure point (**Fig.1**, **Fig. 7**). We first observe elevation of bilateral neural folds to create a keyhole-shaped neural groove, likely driven by Myosin-dependent apical constriction of midline neural plate cells. These folds then pinch together in the center to create anterior and posterior opening that zipper closed away from the pinch point, during which Myosin is enriched at the zipper (**Fig. 1**). The anterior portion of the groove goes on to form the previously described eye-shaped opening (**Fig. 3**)(*29*). The posterior opening has not (to our knowledge) been described before, but a previous time-lapse imaging study of midbrain-hindbrain boundary formation captured zippering of an opening in the hindbrain (*88*) that we speculate is the same posterior closure point. We observed that closure at each point occurs in a reproducible order - first the central pinch point, then the anterior followed by posterior ends of the eye-shaped opening (**Fig. 7**) - raising the possibility that they are analogous to the multiple discrete closure points where neural fold zippering initiates in the mouse (*13, 14*). It was further shown in mice that this closure is facilitated by contact between the non-neural ectoderm (NNE) (*14, 89*), which extends protrusions that meet across the neural groove to “button up” the neural folds (*90*). Our study did not directly address the behavior of NNE and provides no evidence for such protrusions, but neuroectoderm cells were previously shown to extend filopodia and make contact with cells on the contralateral side (*29*). We did observe, however, that the periderm overlying the developing neural tube spanned the neural groove as the neural folds elevated and fused (**Fig. 4**). Whether this thin epithelial sheet contributes to neural tube closure, similarly to the NNE of mouse embryos, will require additional investigation.

### Effect of vangl2 deficiency on neural development

Loss of the planar cell polarity protein and NTD risk gene *vangl2* has dramatic effects on vertebrate neural tube development. While disruption of this gene prevents neural tube closure in mice and frogs (*3, 4, 38, 74*), in zebrafish it produces double neural lumens divided by a mass of neuroectoderm cells that result from abnormal midline C-divisions (*21, 55*). By examining a more anterior region of the developing neural tube in *vangl2* deficient zebrafish embryos, we identified additional phenotypes that more closely resemble those of other vertebrate species. Live imaging revealed that, unlike WTs, *vangl2* deficient neural folds did not zipper smoothly closed away from the pinch point but instead “buttoned” up at a series of ectopic pinch points that emerged at increasingly posterior positions along the neural tube. The openings between these pinch points did eventually close in most *vangl2* deficient embryos examined, but this closure was significantly delayed compared with WT controls. Notably, neural fold fusion itself was not disrupted by loss of *vangl2* (unlike mice (*71*)), and zippering may actually be accelerated in these embryos (**Fig. 3F**). Instead, we speculate that this delay is due to increased width of the neural plate at the time of fusion and therefore, is likely a consequence of reduced CE of neuroectoderm (*21, 27*). Whether delayed and/or abnormal anterior neural fold fusion is a common feature of all mutations that disrupt CE remains to be examined.

In addition to the ectopic “pinching” of the neural folds upon loss of *vangl2*, that these embryos did not exhibit the striking lumen enclosure observed in WTs. Instead, the neural groove of *vangl2* deficient embryos sealed up from ventral to dorsal, leaving a solid neural keel rather than a hollow lumen. Whether and how this phenotype is related to delayed neural fold fusion will be an interesting area for future study. We note that because anterior neural fold fusion is complete (even in most *vangl2* deficient embryos) by the time of onset of C-divisions (*17, 18, 24*), it is unlikely that abnormal C-divisions contribute to the observed defects in forebrain closure. Live imaging also revealed abnormal morphology of the periderm overlying the neural plate in *vangl2* deficient embryos. Although the periderm was also seen spanning the neural folds in WT embryos (**Fig. 4**), its constituent cells were flat. By contrast, *vangl2* deficient periderm cells were rounded and protruded dramatically from the neural groove (**Figs. 2-4**). Whether abnormal periderm cells are contributors to, a consequence of, or unrelated to delayed neural fold fusion in these embryos is unknown.

### Implications for NTD modeling in zebrafish

Zebrafish embryos are highly amenable to genetic and chemical screening techniques (*78, 91, 92*) that enable identification of causative genetic variants and gene-environmental interactions, but their utility in NTD modeling has been limited by their apparent poor resemblance to mammalian neurulation. Researchers have suggested bifurcated pineal precursors in one-day-old embryos as a proxy for NTDs in zebrafish and have even identified gene-environment interactions that exacerbate pineal defects (*30, 32*). However, it was not clear to what extent these phenotypes resemble NTDs because A) they were not examined histologically and the mutations that induce them (in genes encoding Nodal signaling components and N-cadherin) are not associated with human NTDs. Here, we show that loss of an NTD risk gene – *vangl2* - does indeed cause widened and occasionally split pineal domains in the forebrain on day 1 (**Fig. 5**). However, we find that this external phenotype can manifest with a variety of internal phenotypes, some of which show little resemblance to amniote NTDs (**Supp.** Fig. 4). By instead examining anterior neural development at the time of neural tube closure, we avoid the confounding effects of ectopic lumen formation seen at later stages upon loss of *vangl2* or Nodal signaling. Indeed, the delay in neural fold fusion in *tri*-/-mutants and *vangl2* morphants is readily observed within fixed embryos at peri-closure stages, providing an easily screen-able phenotype. However, whether loss of other human NTD risk genes produces similar phenotypes in zebrafish remains to be tested. Given that open neural tubes are only apparent in the forebrain region of zebrafish embryos, it is also not clear whether this is a fitting model for only anterior NTDs (like anencephaly and craniorachischisis) or if mutations causing posterior NTDs like spina bifida would yield similar phenotypes. By providing direct evidence for conservation of fold-and-fuse neurulation within zebrafish, and identifying a readily screen-able phenotype upon loss of an NTD risk gene, our study raises the possibility of using zebrafish to model NTDs in the future.

**Figure 7.**
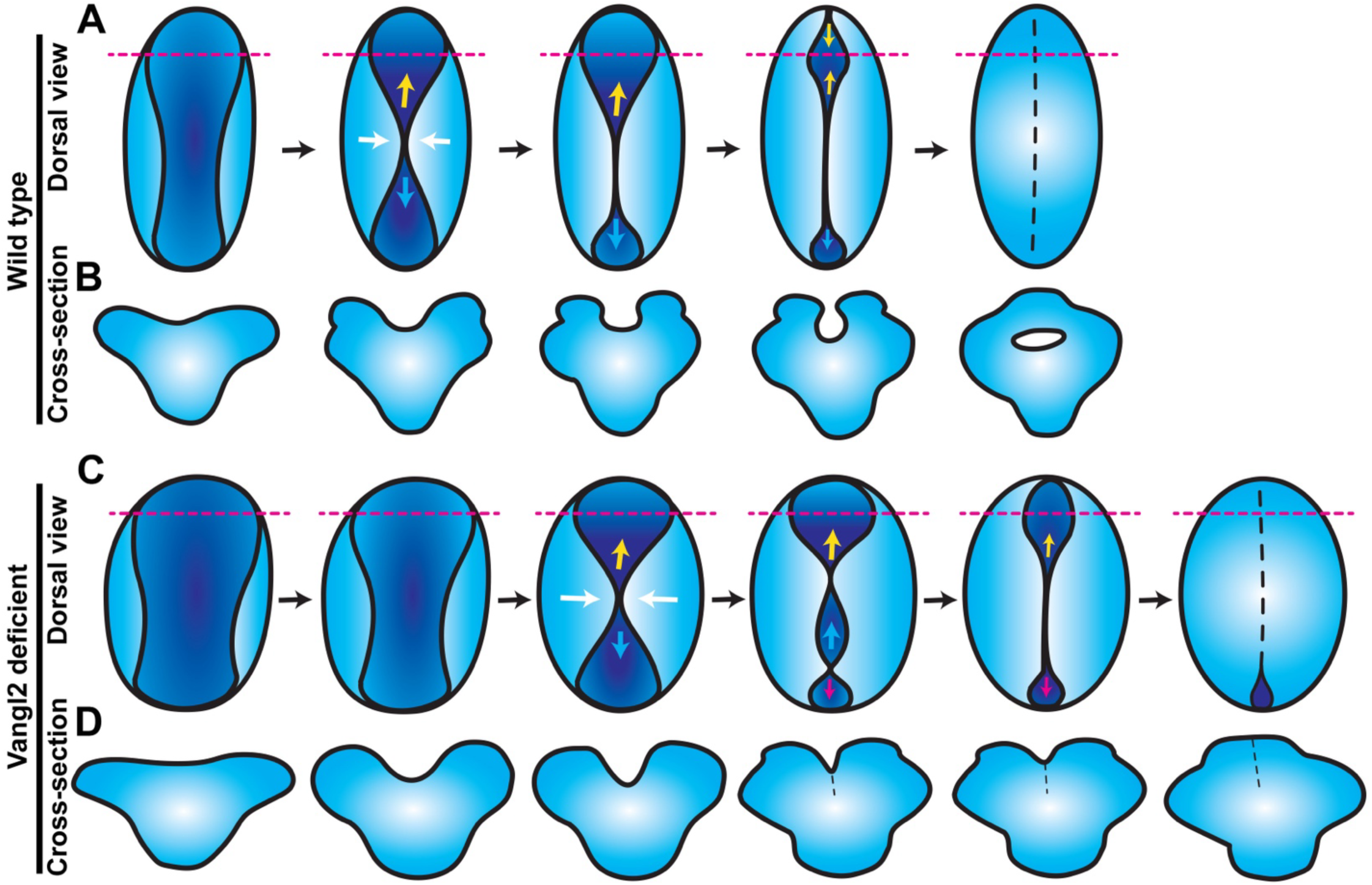
Model for anterior neural tube closure in zebrafish embryos. **A)** Diagram of the anterior (brain region) neural plate in WT zebrafish embryos from approximately 4-10 somite stage, viewed from the dorsal surface with anterior to the top. A shallow neural groove (dark blue) forms at the dorsal midline between the bilateral neural folds (light blue). The neural folds come together at a central “pinch point” (white arrows), creating anterior and posterior openings. The posterior opening zippers closed caudally from the pinch point (cyan arrows) and the anterior opening goes on to form an eye-shaped opening in the forebrain region. The anterior edge of the eye-shaped opening zippers toward the posterior while its posterior edge zippers anteriorly, closing the eye-shaped opening from both sides (yellow arrows). The posterior opening continues to zipper toward the hindbrain until the neural folds in the entire brain region have fused. Dashed magenta lines represent the positions of the cross-sectional views shown in (B). **B**) Cross-sectional views of the anterior WT neural plate at the position of the dashed magenta lines in (A), dorsal is up. A U-shaped neural groove forms between the bilateral neural folds, which approach the midline and then fuse dorsally to enclose a hollow lumen. **C**) Diagram of anterior neural plate morphogenesis in vangl2 deficient embryos. The neural plate and groove begin wider and are delayed in the formation of the first pinch point (white arrows) and neural fold fusion. Additional pinch points form int the posterior opening, creating an additional opening that zippers closed (magenta arrows). **D**) Cross-sectional views of the anterior *vangl2* deficient neural plate at the position of the dashed magenta lines in (C), dorsal is up. The forebrain forms a V-shaped neural groove that seals up from ventral to dorsal rather than enclosing a lumen.

## Materials and Methods

### Zebrafish

Adult zebrafish were maintained through established protocols (*93*) in compliance with the Baylor College of Medicine Institutional Animal Care and Use Committee. Embryos were obtained through natural mating and staging was based on established morphology (*94*). Studies were conducted using AB WT, *tdgf1/oep^tz257^*(*82*), *vangl2*/ *trilobite^vu67^* (*95*), and TgBAC[*flh:flh-kaede*] (*80*) embryos. Fish were crossed from their home tank at random and embryos were chosen for injection and inclusion in experiments at random.

### Microinjection of synthetic mRNA and morpholino oligonucleotides

Single-celled embryos were placed in agarose molds (Adaptive Science tools I-34) and injected with 0.5-2 nL volumes using pulled glass needles (Fisher Sci #50-821-984). mRNAs were transcribed using the SP6 mMessage mMachine kit (Fisher Sci #AM1340) and purified using Biorad Microbiospin columns (Biorad #7326250). Each embryo was injected with 100 pg memGFP or 200 pg mCherry mRNA, and/or 0.5 - 2 ng *vangl2* MO-4 ((*75*) sequence: 5’ - AGTTCCACCTTACTCCTGAGAGAAT - 3’).

### Whole mount in situ hybridization

Antisense riboprobes were transcribed using NEB T7 or T3 RNA polymerase (NEB #M0251s and Fisher #501047499) and labeled with digoxygenin (DIG) NTPs (Sigma/Millipore #11277073910). Whole mount in situ hybridization (WISH) was performed according to (*96*) with minor modifications. Embryos were fixed as described above, washed in PBS + 0.1% Tween-20 (PBT), gradually dehydrated, and stored in methanol at - 20C. Immediately prior to staining, embryos were rehydrated into PBT and hybridized overnight with antisense probes within the wells of a 24-well plate. Embryos were gradually washed into SSC buffer and then into PBT before overnight incubation with an anti-DIG primary antibody at 1:5000 (Roche #11093274910). Embryos were washed in PBT and then staining buffer before developing in BM Purple staining solution (Roche #11442074001). Embryos were washed and stored in stop buffer (10 mM EDTA in PBT) until imaging.

### Histology

After whole mount in situ hybridization, the head regions of 28 hpf embryos were isolated, mounted in Tissue Tek O.C.T. medium (VWR #25608-930) within plastic base molds, and snap frozen in liquid nitrogen. 14 μm serial sections were cut and collected by the Baylor College of Medicine RNA In Situ Hybridization Core.

Sections were mounted under coverslips and imaged using a Nikon Fi3 color camera on the Nikon ECLIPSE Ti2 microscope described below.

### Inhibitor treatments

Nodal inhibitor SB505124 (VWR #103540-834) was stored as a 10 mM stock in DMSO at 4 C. Embryos were dechorionated prior to treatment with 50 μM SB505124 in 0.3x Danieau’s solution at the stages indicated.

Embryos were incubated at 28.5 C within the agarose-coated wells of a 6-well plate until 28 hpf, at which time they were fixed as described above and processed for WISH.

### Phalloidin staining

Embryos were fixed as described above, rinsed in PBT, and stained directly by incubating with Alexa Fluor 546 Phalloidin (ThermoFisher A22283) in PBTr for several hours. Embryos were rinsed in PBTr and mounted for confocal imaging as described below.

### Microscopy

Fixed phalloidin stained embryos were mounted in 3% methylcellulose, and live embryos were mounted in 0.35% low-melt agarose (ThermoFisher #16520100) in glass bottomed 35 mm petri dishes (Fisher Sci #FB0875711YZ) prior to imaging. Confocal Z-stacks were collected using a Nikon ECLIPSE Ti2 confocal microscope equipped with a Yokogawa W1 spinning disk unit, PFS4 camera, and 405/488/561nm lasers (emission filters: 455/50, 525/36, 605/52). Confocal Z-stacks were obtained with a 1 μm (fixed embryos) or 2 μm (live embryos) step size using a Plan Apo Lambda 20X lens. For time-lapse series, embryos were maintained at 28.5 C in a Tokai Hit STX stage top incubator and Z-stacks were collected at 5-minute intervals. Images of WISH-stained embryos were taken with a Nikon Fi3 color camera on a Nikon SMZ745T stereoscope.

### Image analysis

ImageJ/FIJI was used to visualize and measure all microscopy data sets. Researchers were blinded to the conditions of all image data using the *blind_renamer* Perl script (https://github.com/jimsalterjrs/blindanalysis?tab=readme-ov-file) (*97*) prior to analysis. Measurements of embryonic structures from fixed embryos, including pineal precursors and somites, were made by drawing a line from one side of the structure to the other at its widest point. Distance between neural folds, neural plate width, and neural grove angle and cross-sectional area were measured similarly from images of live embryos. 3D projections of confocal z-stacks were made using the ‘3D project’ plugin.

### Statistical Analysis

Graphpad Prism 10 software was used to perform statistical analyses and to generate graphs for the data collected during image analysis. Datasets were tested for normality prior to analysis and statistical tests were chosen accordingly. The statistical tests used for each data set are noted in figure legends.

## Acknowledgements

We thank Dr. Patrick Blader for generously sharing the Tg[*flh:kaede*] fish line, Dr. Lila Solnica-Krezel for sharing additional fish lines and plasmids, and Dr. Lance Davidson for sharing the sf9-mNeon plasmid. Histology services were provided by Dr. Cecilia Ljungberg and Rong Jyh Kao of the BCM RNA In Situ Hybridization Core and animal care was provided by the BCM Center for Comparative Medicine. We thank Drs. Rachel Brewster, Dan Gorelick, and Ryan Gray for helpful discussions and comments on the manuscript. Thanks to Williams lab members for technical assistance, support, and feedback on this project.

## Supplemental video legends

**Supp. video 1**: Posterior view of neural groove formation in the forebrain region of a wild-type zebrafish embryo labeled with membrane-GFP. Movie begins at approximately the 4 somite stage and each frame = 5 minutes. Video shows a single Z plane of a 3D confocal time series.

**Supp. video 2**: Anterior view of neural fold fusion in the forebrain region of a wild-type zebrafish embryo labeled with membrane-GFP. Movie begins at the 4-5 somite stage and each frame = 5 minutes. Video shows a single Z plane of a 3D confocal time series.

**Supp. video 3**: Anterior view of neural groove formation in the forebrain region of a wild-type zebrafish embryo labeled with membrane-mCherry and the Sf9-mNeon Myosin reporter. Movie begins at approximately 3 somite stage and each frame = 5 minutes. The left and right panels show a single Z plane at deeper and more superficial positions in the Z-stack, respectively.

**Supp. video 4**: Anterior view of neural fold fusion in the forebrain region of a wild-type zebrafish embryo labeled membrane-mCherry and the Sf9-mNeon Myosin reporter. Movie begins at the 4-5 somite stage and each frame = 5 minutes. Video shows a single Z plane of a 3D confocal time series.

**Supp. video 5**: Neural fold fusion in the forebrain region of a wild-type zebrafish embryo labeled with membrane-GFP. Movie begins at the 6-7 somite stage and each frame = 5 minutes. Video shows a single Z plane of a 3D confocal time series.

**Supp. video 6**: Posterior view of neural groove formation in the forebrain region of a *tri*-/- mutant zebrafish embryo labeled with membrane-mCherry. Movie begins at the 6-7 stage and each frame = 5 minutes. Video shows a single Z plane of a 3D confocal time series.

**Supp. video 7**: Posterior view of neural fold fusion in the forebrain region of a *tri*-/- mutant zebrafish embryo labeled membrane-mCherry and the Sf9-mNeon Myosin reporter. Movie begins at the 6-7 somite stage and each frame = 5 minutes. Video shows a single Z plane of a 3D confocal time series.

**Supp. video 8**: Neural fold fusion in the forebrain region of a *vangl2* morphant zebrafish embryo labeled with membrane-GFP. Movie begins at the 6-7 somite stage and each frame = 5 minutes. Video shows a single Z plane of a 3D confocal time series.

**Supp. video 9**: Optical transverse section through the posterior forebrain region of a wild-type zebrafish embryo labeled with membrane-GFP. Movie begins at the 4 somite stage and each frame = 5 minutes. Video shows a single Z plane of a 3D confocal time series.

**Supp. video 10**: Optical transverse section through the posterior forebrain region of a *tri*-/- mutant zebrafish embryo labeled with membrane-GFP. Movie begins at the 4 somite stage and each frame = 5 minutes. Video shows a single Z plane of a 3D confocal time series.

**Supp. video 11**: Optical transverse section through the posterior forebrain region of a *vangl2* morphant zebrafish embryo labeled with membrane-GFP. Movie begins at the 4 somite stage and each frame = 5 minutes. Video shows a single Z plane of a 3D confocal time series.

